# The plant circadian clock exerts stronger control over the diel proteome than the transcriptome

**DOI:** 10.64898/2025.12.19.695194

**Authors:** Devang Mehta, Mohana Talasila, Zhi Xing Lau, Maria Camila Rodriguez Gallo, Qiaomu Li, Yangyi Zhong, Sukalp Muzumdar, Roy Li, Wei Jin Luo, Vincent Lau, Asher Pasha, Sarah Lock, Daphne Ezer, Nicholas Provart, R. Glen Uhrig

## Abstract

The plant circadian clock is a genetic circuit composed of multiple mutually-regulating transcription factors that together synchronize internal biological rhythms to the ∼24-hour period of planetary rotation. While it has been known for over a decade that nearly 40% of the transcriptome in the model plant Arabidopsis oscillates with a circadian rhythm, it is yet unclear to what extent this translates to the proteome. Here, through parallel quantitative proteome and transcriptome time-course profiling of Arabidopsis wild-type plants and a panel of clock deficient plant lines, we show that specific clock genes exercise extensive control over diel proteome rhythmicity, and to a much greater extent than they do the transcriptome. This control results in a clock-dependent synchronization of rhythmic proteins along a bimodal phase distribution that is lost in circadian clock deficient plants. This suggests pervasive post-translational control of gene expression by specific elements of the circadian system, notably the morning expressed LHY/CCA1 module. Our findings imply that the circadian clock exercises much greater control of gene expression through proteostasis mechanisms than previously recognised, necessitating a recalibration of our current understanding of clock proteins as primarily transcriptional regulators.

## Main Text

Most eukaryotic species, and some bacteria, maintain a circadian clock to time growth, development and metabolism to daily fluctuations in external stimuli timed in accordance with the rotation of the planet(*1*, *2*). However, as sessile organisms beholden to the presence of light to drive growth and development, plants have independently evolved a complex clock network comprised of multiple interconnected transcriptional regulators(*3*). With its far-reaching regulatory capacity, this network both directly and indirectly controls nearly all physiological processes in plants, from responses to abiotic and biotic stress to seasonality and life cycle(*4–7*).

To date, much of what we know about the plant circadian clock has been driven by a combination of genetics and transcriptomic experimentation under primarily free-running conditions using the model plant Arabidopsis. In free-running experiments, plants are first entrained by growth under light-dark cycles followed by growth under continuous light or dark for circadian measurement. This is done to distinguish between the impact of light signalling and the endogenous biological oscillations maintained by the clock network. These pioneering studies have paved the way for our current understanding of the plant circadian clock as a network of transcriptional regulators acting primarily through their binding and regulation of target promoter elements, in both other clock genes, and those comprising downstream pathways such as growth and development. This transcription-focused understanding of plant clock function has been reinforced by time-course transcriptomics experiments, which have consistently found between 30-40% of transcripts oscillate with a circadian rhythm(*8*). The maintenance of these rhythms under free-running cycles is often cited as evidence that a large proportion of the plant transcriptome operates under circadian control(*9–11*). Furthermore, previous research examining circadian changes in the plant proteome has found limited evidence of diel protein abundance changes(*12–17*), supporting a primarily transcriptional clock.

On the other hand, however, chromatin immunoprecipitation sequencing (ChIP-Seq) experiments with individual circadian clock transcription factors has routinely discovered fewer direct targets—on the order of hundreds, rather than the thousands of oscillating transcripts—suggesting a gap in our current model of gene expression regulation by the clock(*18–26*). Targeted biochemical research has also found associations between clock proteins and proteasomal regulation, most prominently through the clock-associated ZEITLUPE (ZTL) protein(*27*, *28*). Further, advances in mass spectrometry technologies and quantitative proteomic data acquisition workflows now suggest that the previously observed lack of circadian clock control of proteome abundance might have been due to technical limitations caused by a strong bias towards high-abundant proteins in past workflows(*29*). For instance, more recent work using the newer BoxCarDIA proteomics data acquisition method has revealed a greater role for diel proteasomal regulation by clock genes(*30*).

Hence, we aimed here to disentangle the relative contribution of the plant circadian clock to gene expression regulation at the transcript and protein levels using a newer, less biased mass-spectrometry based proteomics methodology together with parallel RNA-seq based transcriptomics. In contrast to earlier studies, we sought to directly estimate the circadian clock’s contribution to diel gene expression regulation by simultaneously profiling a panel of clock mutants under equinox growth conditions rather than focusing on wild-type plants under free-running cycles. Here we find that a greater proportion of the proteome oscillates with a diel rhythm than detected in past studies. Our parallel proteome-transcriptome analysis also showed that the circadian clock genes have a far greater impact on diel proteome oscillations than diel transcriptome oscillations, likely through the dysregulation of proteostatic mechanisms. In addition to new insights into diel and circadian control of biological processes at the protein-level, our results provide a fundamentally new understanding of how, and to what extent, the plant circadian clock network acts post-transcriptionally to control plant biology.

## Clock genes control the bimodal phasing of the 24-hour rhythmic Arabidopsis proteome to varying degrees

We harvested whole rosette tissue from 24-day old soil-grown Arabidopsis plants, including wild-type (WT) and six well-characterized clock-deficient plant lines encompassing key circadian transcriptional regulators expressed at different times throughout the day (*lhy-20 cca1-1, prr7-3 prr9-1, prr5-11 prr7-11, toc1-2, elf4-101* and *lux-4*). Mature, pre-bolting rosettes were harvested every two hours over a 24-hour period under a 12:12 equinox light regime, with four independent pools of 3 plants harvested per genotype per sample at each timepoint (n=4). In order to obtain uniform and comprehensive proteomes for each plant sample, we deployed our recently developed BoxCarDIA methodology for liquid chromatography mass-spectrometry (LC-MS) with label-free quantitation using the library-free data independent acquisition (DIA)(*29*, *31*). This resulted in a final dataset of 332 high-quality proteomes. We quantified a median of 6,445 protein groups from 24,496 peptides per sample, with a minimum of 4,800 protein groups quantified across all samples; a substantial improvement over past time-course proteome datasets in plants (Fig. 1A & B; Data S1).

**Fig. 1.**
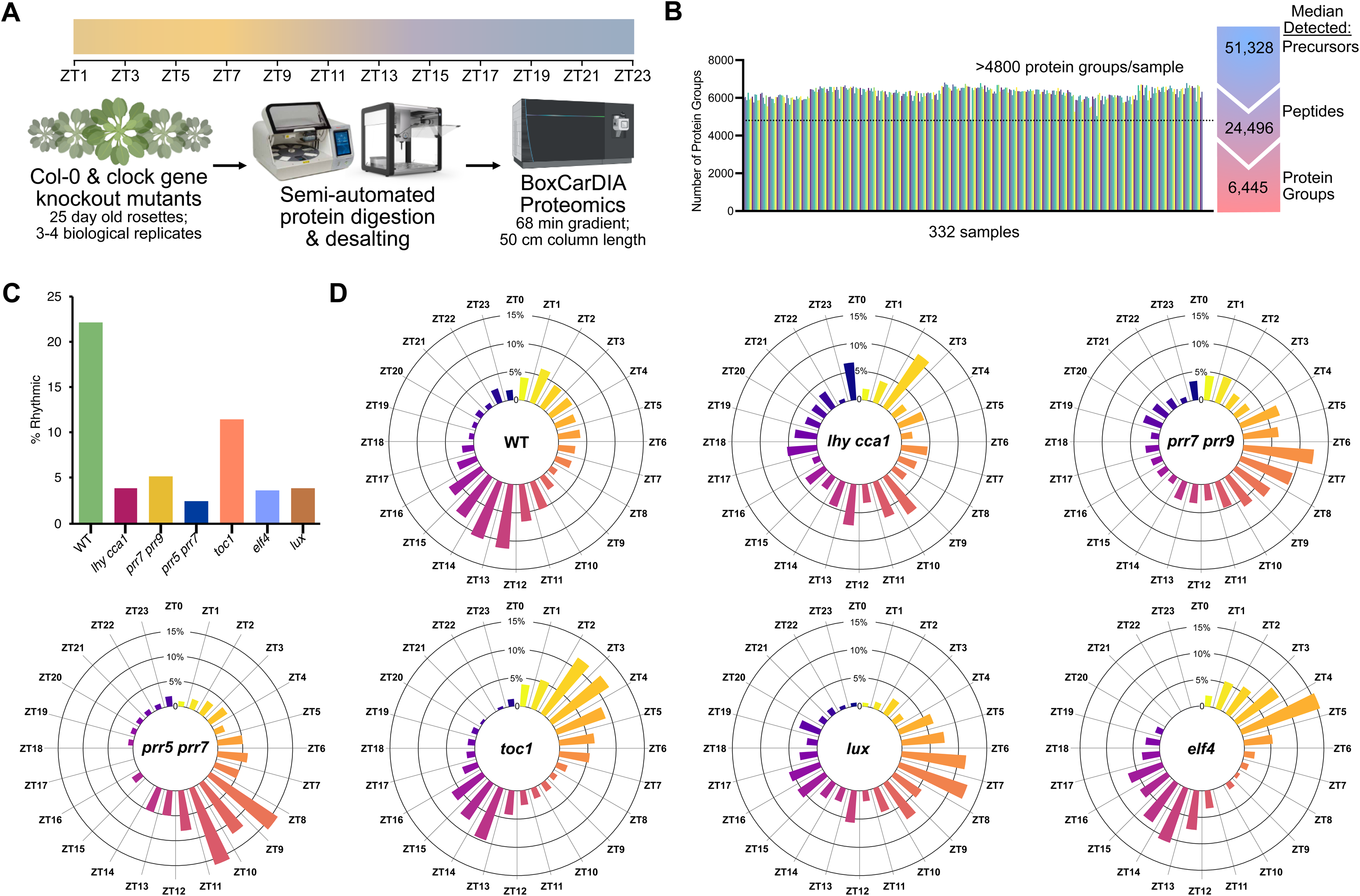
**The synchronisation of the plant diel proteome and its control by circadian clock genes**. Wild-type (WT) *Arabidopsis thaliana* Col-0 rosettes were processed using a semi-automated workflow and analysed by quantitative proteomics (*see materials and methods*). (A) Visual schematic of the quantitative proteomic workflow and the resulting median quantified precursors, peptides. (B) Protein groups quantified in each of the 332 analysed samples and median protein-groups per sample across the dataset (Data S1). (C) Percent rhythmic proteins in WT and six different circadian clock deficient plant lines (Cosinor; Data S2). (D) Radial plots of peak time of day (zeitgeber time; ZT) abundance for each significantly oscillating protein in WT and six circadian clock deficient plant lines.

Next, to discover the proportion of the proteome that oscillated with a circadian rhythm in all genotypes, we performed rhythmicity analysis using the JTK-Cycle and Cosinor algorithms implemented in the DiscoRhythm software package(*32*). Through this analysis, we estimated that 22% of the quantified proteins in wild-type Arabidopsis plants exhibit diel oscillations in their abundance over the course of a day. This is markedly higher than previous data-dependent acquisition (DDA) proteomics studies which found that only between 3–9% of the Arabidopsis proteome exhibited diel rhythmicity, after reanalysis with the same parameters (*13*, *15*). In contrast to wild-type plants, only 2.5–5% of each clock mutant proteome exhibited diel rhythmicity, excepting the weak mutant *toc1*, which showed a 10% proteome-level rhythmicity (Fig. 1C; Data S2). In order to validate that these differences are truly due to differential phasing of the rhythmic proteome between wild-type plants and mutants, and not global differences in protein abundance, we also performed daily expression integral (DEI) analysis. Here, we calculated the daily integral of the abundance of each protein and compared this for each mutant against the wild-type data (fig. S1A). For each mutant, the proteome-level DEI correlated to a remarkable degree to wild-type, affirming that rhythmicity differences observed are not due to any systemic differences in protein abundance between genotypes. The notably lower rhythmicity of the proteome of circadian clock mutants suggests that the clock plays a greater role than previously recognised in maintaining rhythmic gene expression patterns under standard light-dark cycles.

We next asked if rhythmic proteins in wild-type plants are synchronized to oscillate with the same phase, i.e., do their abundances peak and fall at similar times of day? To investigate this, we plotted the time-of-day of peak abundance (acrophase) of proteins detected as rhythmic in wild-type plants and each circadian clock deficient plant line profiled here (Fig. 1D; Data S2). We observed that rhythmic proteins in wild-type plants are phased along a bimodal distribution with a large population of proteins synchronized to reach their peak abundance at the onset of night (ZT 11-16) and a second, smaller, population of proteins peaking throughout the day. Curiously, it appeared that very few proteins (<10%) reached peak abundance from the middle of the night to the start of the day (ZT 18-23). This bimodal pattern was highly disrupted in the *lhy cca1* double mutant, demonstrating for the first time that the clock controls the synchronicity of rhythmic proteins. Rhythmic protein phasing, however, was different in each clock deficient plant line, with some (e.g. *toc1* and *elf4*) appearing closer to wild-type (albeit with more morning-phased proteins) and others (e.g. *prr7 prr9* and *prr5 prr7)* demonstrating more unimodal protein synchronicity. Interestingly, the acrophase profile of *lux* was notably different from *elf4*, with proteins peaking later in the day and night. LUX is the key DNA-binding component of the clock Evening Complex together with ELF4 and ELF3 (which lack DNA-binding capacity)(*33*, *34*). The different acrophase profiles of *lux* and *elf4* suggests that the two genes have Evening Complex-independent roles in controlling rhythmic proteome synchronization. In addition to the phase of rhythmic abundance, we also assessed if there were global changes in the amplitude of rhythmic proteins (fig. S2A). Here we found that similar to the results on overall rhythmicity, the rhythmic proteome of *lhy cca1* showed lower amplitude, while that of *toc1* oscillated with similar amplitude as the wild-type plants, reflecting the greater rhythmicity and weaker phenotype associated with this mutant.

## Clock genes control diel rhythmic gene expression post-translationally

To further contextualize our proteome-level results, we acquired complementary RNA-seq data from the very same rosette tissue for the WT, *lhy cca1, prr7 prr9* and *toc 1* genotypes (Fig. 2A; Data S3). These mutants were selected to profile the transcriptome of plants that display a range of different proteome rhythmicity phenotypes: strong asynchronization (*lhy cca1*), unimodal synchronization *(prr7 prr9*), and wild-type like bimodal synchronization (*toc1*). Of the 192 total samples, we obtained 187 transcriptomes that passed our quality criteria, with a minimum of 3 replicates per genotype and per timepoint. Principal Component Analysis of the transcriptome data revealed clear clustering of samples by time-point across all genotypes, with limited overall transcriptome differences between each genotype (fig. S3; Data S4). By contrast, proteome data clearly clustered by genotype to a greater extent (fig. S3; Data S3). As an initial benchmark check, we checked that gene expression profiles for clock genes in our wild-type transcriptomes tallied with published data, finding a high-level of concordance with previously published diel microarray datasets (fig. S4)(*35–37*).

**Fig. 2.**
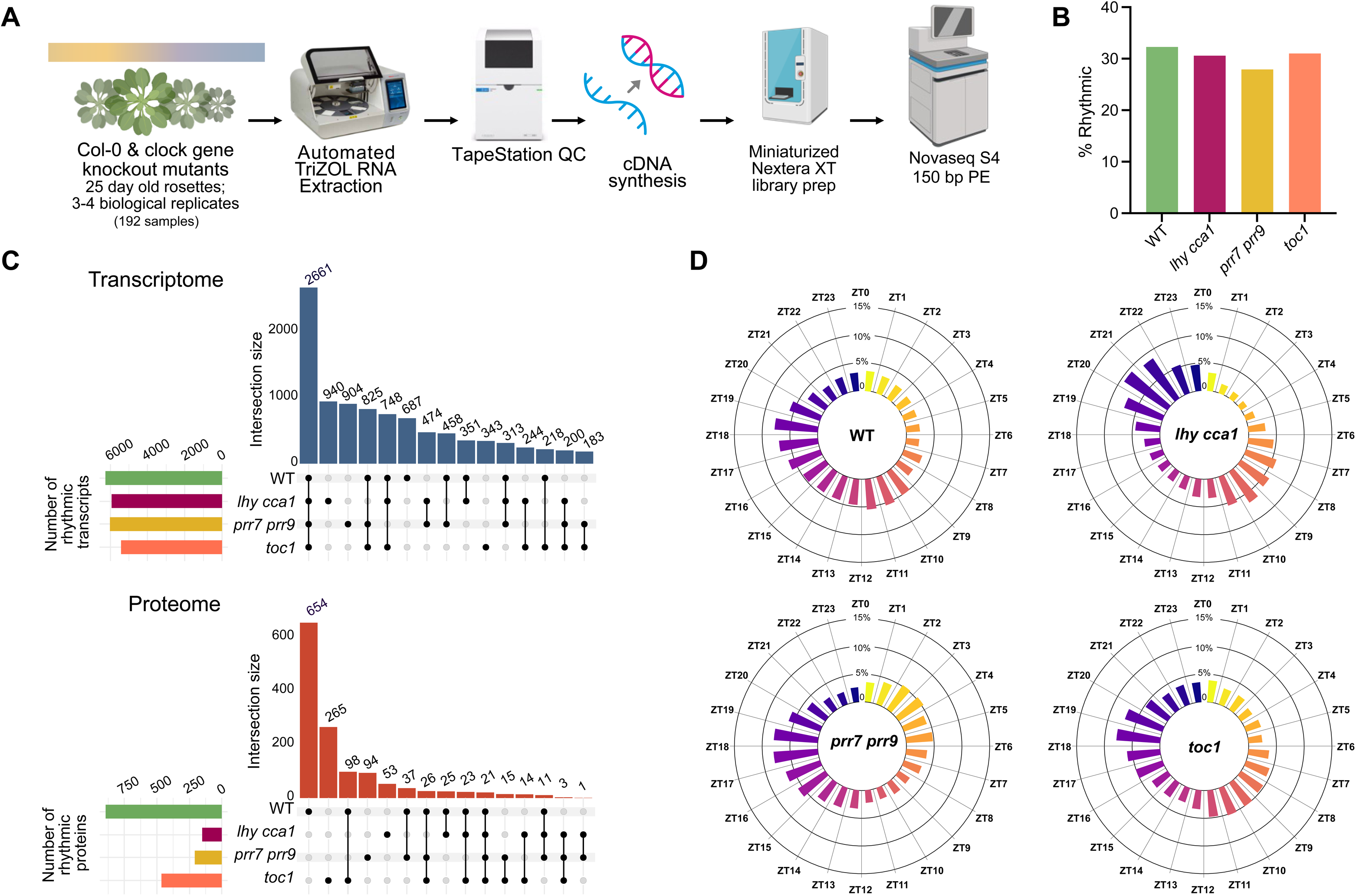
**Circadian clock genes have a limited impact on the Arabidopsis diel transcriptome**. RNAseq analysis of the same samples subjected to quantitative proteomic analysis was performed using the genotypes WT, *lhy cca1*, *prr7 prr9* and *toc1* (Data S3). (A) Schematic of the RNAseq workflow. (B) Percent rhythmic transcripts in WT and the three circadian clock deficient plant lines analysed (Cosinor; Data S4). (C) Upset plot of the cycling transcriptome and proteome across each examined genotype. (D) Radial plots depicting peak time-of-day (zeitgeber time; ZT) abundance for each significantly oscillating transcript in WT and the three circadian clock deficient plant lines analysed.

Next, normalized read counts from this dataset were used for rhythmicity analysis, applying highly stringent FDR thresholds and only classifying genes detected as rhythmic by both Cosinor and JTK-Cycle as rhythmic transcripts. Unsurprisingly, 32.29% of the wild-type transcriptome oscillated with a ∼24 h period, consistent with previous microarray and RNA-seq datasets(*36*, *37*). However, to our surprise, and unlike our proteome data (Fig. 1C), a similar proportion of the transcriptome was rhythmic in the three different clock mutants profiled here (30.58%, 27.9% and 31.02% in *lhy cca1*, *prr7 prr9,* and *toc1*, respectively) (Fig. 2B; Data S6). Similar to our proteome data, we also performed a DEI correlation analysis of transcript levels between each mutant and wild-type. Similar to proteome data, we find a high degree of correlation in overall daily transcript abundance between the mutants and wild-type, suggesting no systemic regulation of abundance (as opposed to phasing) by the clock (fig. S1). Furthermore, analysis of the amplitudes of rhythmic transcripts between genotypes revealed no differences (unlike at the protein level) (fig. S2B), reinforcing the finding that clock mutants have greater disruption in rhythmicity at the protein level than at the transcript level.

Comparing the rhythmic transcripts and proteins detected in each genotype revealed interesting differences between genes specifically altered in different clock mutants (Fig. 2C). These genes, which are rhythmic exclusively in different clock mutants, are genes with arrhythmic expression in wild-type plants and hence likely have highly light-regulated gene expression that is buffered or modulated by individual parts of the clock. At the transcript level, there was considerable overlap between transcripts that retained their diel rhythmicity in *toc1* and wild-type plants, suggesting that *TOC1* does not substantially affect diel gene expression. Intriguingly however, 265 proteins were rhythmic in *toc1* alone, suggesting that the abundance of these proteins is highly light-regulated, but that TOC1 plays a role in buffering their light sensitivity to keep them arrhythmic in wild-type plants (Fig. 2C; fig S5; Data S6). Several of these *toc1* rhythmic genes are linked to carbohydrate metabolism, reinforcing recent research that suggests *TOC1* functions as a rheostat to regulate metabolism under diel cycles(*38*). By contrast, while nearly 940 transcripts were diel rhythmic exclusively in *lhy cca1*, only 53 proteins were diel rhythmic exclusively in *lhy cca1*. This suggests that *LHY / CCA1* plays a relatively smaller role in specifically buffering the diel expression of light-regulated genes in wild-type plants at the protein level but does play a major role in this process at the transcript-level (Fig. 2C; fig S6; Data S6).

Proteomic data discussed above showed a high degree of phase synchronisation regulated by different clock components, most prominently, *LHY / CCA1*. A similar analysis was performed on the rhythmic transcriptome to assess to what degree phase synchronisation is a proteome-specific phenotype. Our results showed that the rhythmic transcriptome of wild-type plants was differently phase synchronised compared to the proteome, with a majority of rhythmic transcripts peaking at the middle of the night (ZT 16-20) (Fig. 2D). Fascinatingly, the rhythmic transcriptome of *lhy cca1* plants showed a bimodal phase distribution reminiscent of the wild-type proteome. This can be partially explained by the fact that many of these transcripts are non-oscillatory in wild-type plants, and whose expression is likely overly light-regulated without the presence of a robustly functional clock in *lhy cca1* plants. Overall, while the extent of rhythmicity observed in plant proteins and transcripts was far greater than that previously observed in mammalian tissue, differential phase synchronization of the transcriptome and proteome in plants was found to correspond well to patterns previously observed in mammalian livers and the fungi, *Neurospora crassa* (*39*, *40*). This suggests an intriguingly conserved diel synchronicity in gene expression in evolutionary distant taxa that have independently evolved circadian clocks.

## Diel compartmentalisation of plant biology at the protein level

Having established that up to 22% of the quantified diel proteome of wild-type plants oscillates with a 24-h period, we next sought to map the biological functions of these rhythmic proteins using functional association analysis involving the String(*41*) and SUBA(*42*) databases (Fig. 3; fig S7). This revealed diversity in terms of temporal compartmentalisation of protein abundances within different biological processes. As expected, primary metabolic processes such as sugar metabolism and photosynthesis are highly compartmentalised by time-of-day. Interestingly however, specialized metabolic processes such as glucosinolate and glutathione metabolism are highly temporally constrained, with each detected enzyme peaking at similar times of the day (i.e., in the dark). We also discovered a previously unrecognised diversity in time-of-day abundance of proteins involved in the metabolism of different amino acids: chloroplast-synthesised branched chain amino acids in the morning, and tyrosine in the evening, for example. Our analysis also revealed intriguing divergence in cytosolic and plastidial translation, with all detected components of the cytosolic ribosome found to peak in the evening. On the other hand, chloroplast encoded ribosomal proteins were found to peak in the morning, while those encoded by nuclear genes peaked in the evening, together with cytosolic ribosome components. This divergence in the temporal regulation of plastidial ribosome components has, to the best of our knowledge, not been discovered previously and warrants further targeted investigation (Fig. 3).

**Fig. 3.**
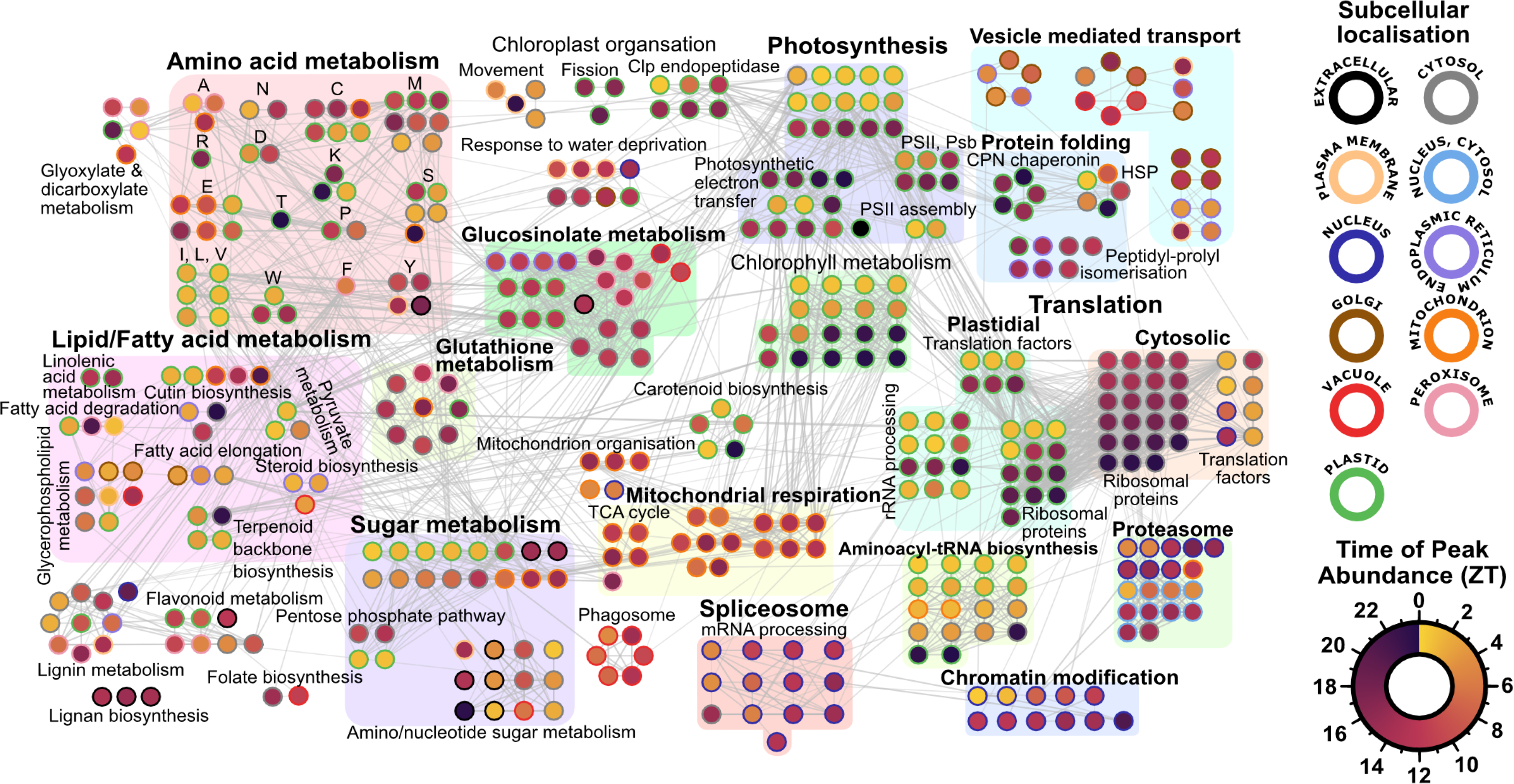
**The functional-temporal partitioning of the diel rhythmic Arabidopsis proteome**. A functional association network analysis assessing the interconnectedness of genes exhibiting a significant change in diel proteomic abundance (Cosinor; Data S2). Node outlines represent the subcellular localization of that protein (SUBAcon 5.0), while node shading depicts the zeitgeber time (ZT) of peak protein abundance. Genes with cycling changes in protein abundance that do not have a STRING-DB edge score > 0.7 to another node are not depicted. Nodes are clustered based on biological process annotation (STRING-DB; https://string-db.org/).

## The role of the clock in coordinating transcriptome and proteome diel oscillations

Since we now had both diel rhythmic transcriptome and proteome datasets for wild-type plants and 3 clock deficient plant lines (*lhy cca1*, *prr7 prr9* & *toc1*), we were able to investigate time-of- day differences, i.e. the phase delay (ΔZT), between transcript and protein oscillations (Fig. 4A). Here, in wild type plants, we were able to observe that for most genes, the phase delay between oscillating transcripts and proteins was within 6 hours, with smaller groups of proteins exhibiting a stronger delay of 11-12 hours and ∼18 hours. In general, the phase delay between rhythmic transcripts and proteins appears to be shorter in plants than similar analysis done in human liver tissue(*39*). While the phase delay pattern was overall similar to wild-type in *prr7 prr9* and *toc1* plants, all the mutants exhibited a shift towards longer phase delays, exemplified by the striking fraction of genes with a phase delay of >18 hours between the transcriptome and proteome in *lhy cca1*, particularly (Fig. 4A). This further reinforces a potential role for the *LHY / CCA1* module in maintaining proteome rhythmicity under diel conditions.

**Fig. 4.**
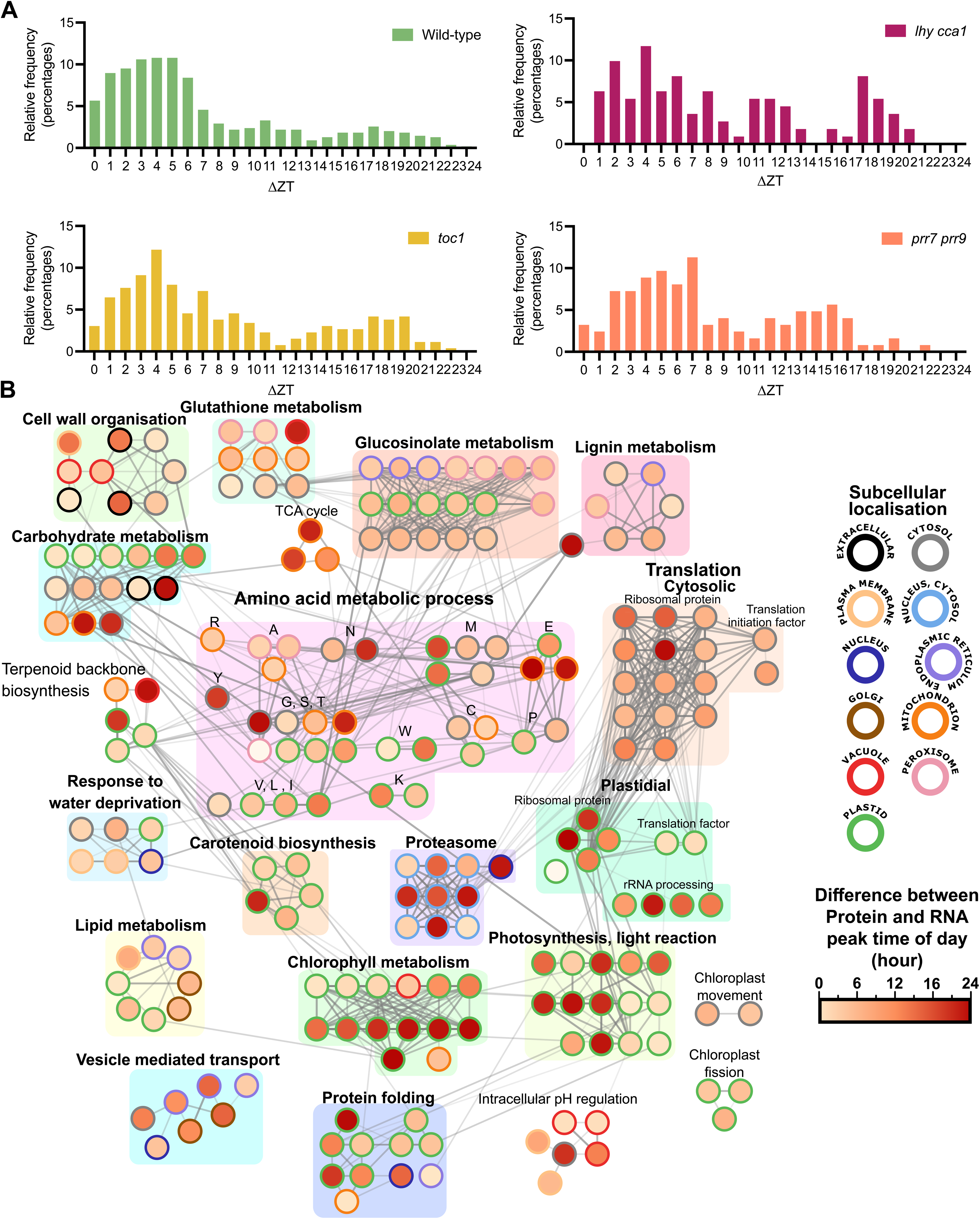
Phase differences between the diel rhythmic transcriptome and proteome. (A) Distribution of the phase difference (absolute difference in peak transcript and protein time-of-day, ΔZT) for each gene with both cycling transcripts and protein abundance. Genotype specific distributions for WT, *lhy cca1*, *prr7 prr9* and *toc1* are shown. (B) Functional association network analysis (STRING-DB) assessing the interconnectedness of genes exhibiting a significant diel change in both transcriptional and proteomic abundance (Cosinor; Data S2, S4 & S5). Node outlines represent the subcellular localization of that protein (SUBAcon 5.0), while node shading depicts delay (ΔZT) between peak transcript and protein abundance. Genes peaking in either transcript or protein abundance only or that do not have a STRING-DB edge score > 0.7 are not depicted. Nodes are clustered based on biological process annotation (STRING-DB; https://string-db.org/).

We next used a similar functional association network analysis to map the phase delay between transcript and protein rhythms for wild-type plants (Fig. 4B). Intriguingly, we found similarities between our results here and our analysis of protein phasing (Fig. 3). Components of processes such as cytosolic translation, carbohydrate metabolism, photosynthesis that had more diel distributed protein phasing also exhibited a wide phase delay range. On the other hand, processes such as glucosinolate metabolism and glutathione metabolism, where proteins were highly synchronized in their peak time–of-day, also had extremely limited phase delay between transcript and protein peak ZTs. This suggests that for such specialized metabolic pathways, concordant regulation of diel rhythms at both the transcript and protein levels is the norm, while other processes are more separately regulated at the transcript and protein levels.

## Clock transcription factors impact the temporal expression of proteostatic genes

Plausible mechanisms explaining the fact that the phase delay pattern between protein and transcript oscillations is impacted by the clock (particularly *LHY / CCA1*) are (a) clock control of protein degradation through the proteasome, or (b) clock-control of translation. Hence, we next compared how diel rhythmic proteins involved in these processes are altered at the protein and transcript levels in the clock deficient plants *lhy cca1*, *prr7 prr9,* and *toc1* (Fig. 5; fig S8; Data S7). Here, we hypothesized that if clock control of proteostatic processes is critical to regulate the phase delay between transcripts and proteins, we should observe a lack of oscillation of proteostasis genes at the protein level in clock mutants, while simultaneously maintaining more wild-type like RNA-level oscillations, which are more influenced by light entrainment (reflecting our finding in Fig. 2B). Indeed, this is what we observe. Both cytosolic and plastidial translation machinery, along with the proteasome, exhibit extensive diel protein-level oscillations that are completely lost in the circadian clock mutants, while corresponding changes in the transcriptome remain largely consistent with WT plants (Fig. 5; Data S7).

**Fig. 5.**
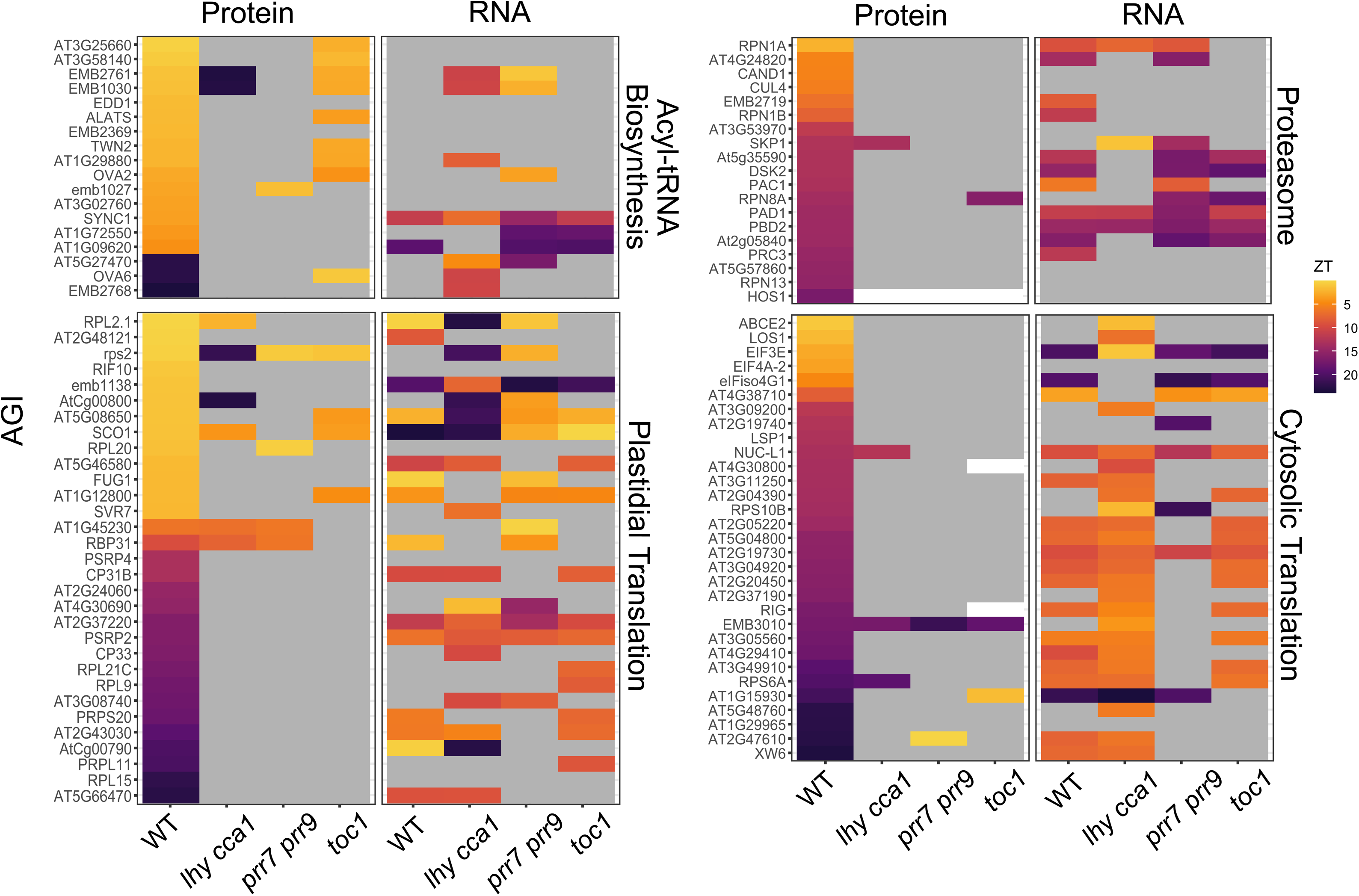
Diel regulation of proteostatic genes is clock-dependent at the protein level. Genes involved in proteostasis-related plant cell processes: acyl-tRNA biosynthesis, plastidial translation, cytosolic translation and proteasomal protein degradation, whose proteins are diel rhythmic on our analysis are heat-mapped based on their peak ZT at both protein and RNA-levels in wild-type (WT) *Arabidopsis thaliana* and *lhy cca1*, *prr7 prr9* and *toc1* lines (Data S7). Grey represent transcripts and proteins that were quantified but not oscillating, and white represents those that were not detected.

To study translational control of rhythmic proteins further, we contextualised our WT data with a recently published ribosome-sequencing (Ribo-Seq) dataset from Arabidopsis plants grown under equinox diel conditions(*43*). To do this, we subset the 895 rhythmic proteins detected in our proteome data based on whether they were also found to be rhythmically regulated in ribosome profiling data (Fig. 6A; Data S8-S9). Through this process, we triaged genes based on (i) rhythmicity and (ii) phasing of their levels at the protein, ribosome-bound, and transcript-level (within 3-hour windows), and classified genes as likely transcriptionally, translationally or post-translationally -regulated, or regulated at multiple levels (Fig. 6A; Data S8-S9). For instance, 207 rhythmic proteins, belonging to the biological processes of aminoacyl tRNA biosynthesis and amino acid biosynthesis, among others, were arrhythmic in the ribosome profiling data, despite cycling in our proteome data. Of the 688 rhythmic proteins that were also found to be rhythmic in the Ribo-Seq data, 562, were rhythmic with a very different phase, and of these, 184 proteins were found to oscillate with the same phase in our RNA-seq and the Ribo-Seq datasets, and a different phase at the protein level. Hence, the oscillatory expression of these 391 (207 + 184) proteins, comprising several tRNA biosynthesis and amino acid biosynthesis genes, is likely caused solely through protein-level regulation. Through similar reasoning, we were able to classify 195 oscillating proteins as likely controlled through both translation and post-translational processes, 110 as controlled transcriptionally, translationally, and post-translationally, 107 as predominantly translationally controlled, 29 regulated at transcription and translation, and only 63 as exclusively transcriptionally controlled (Fig. 6A; Data S8-S9).

**Fig. 6.**
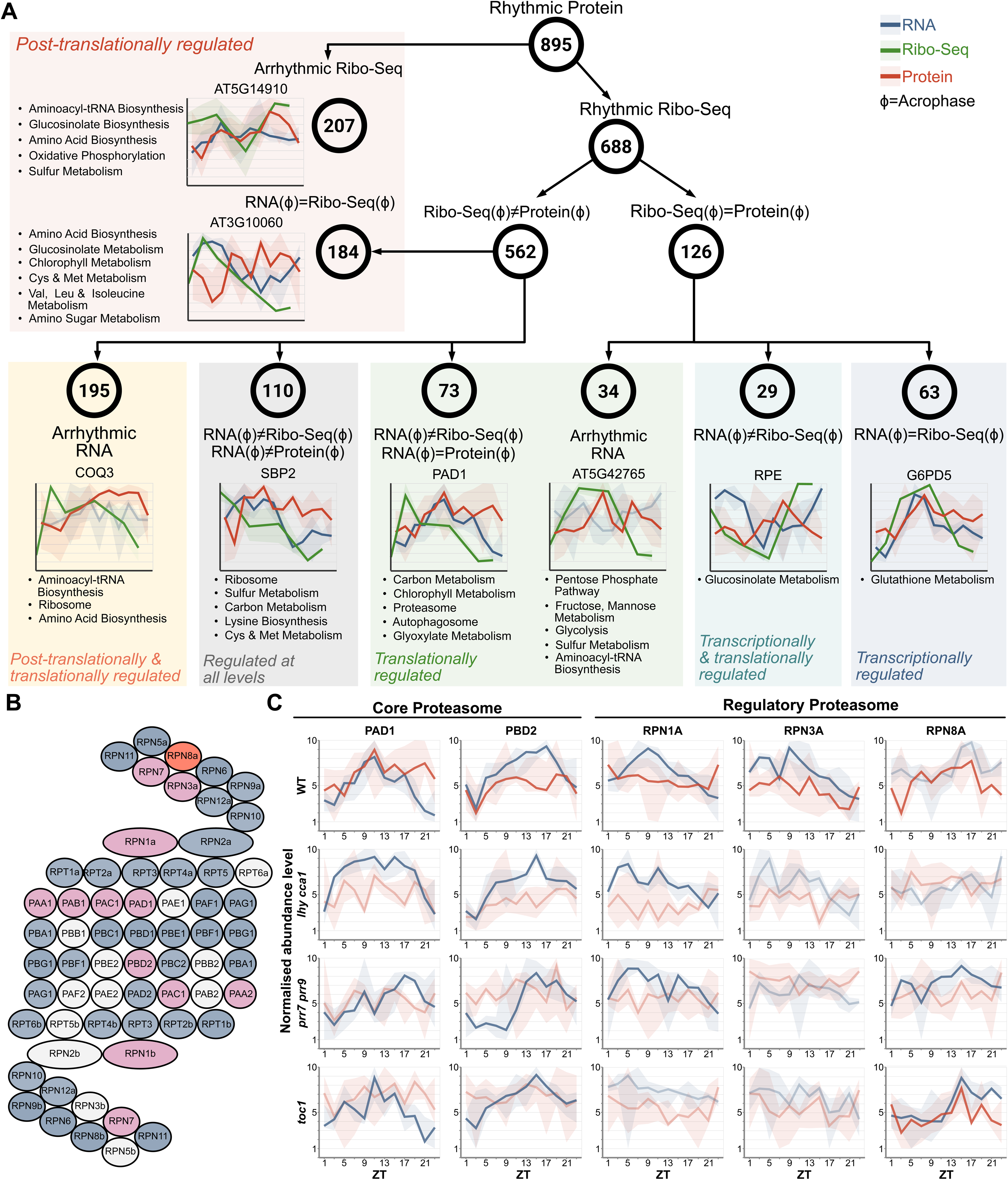
Partitioning of rhythmic proteins in Arabidopsis by likely mode of diel expression regulation based on comparative analysis of proteomics, RNA-seq, and Ribo-Seq datasets. (A). The 895 rhythmic proteins in our dataset were partitioned based on overlap between rhythmicity and phase with their transcripts and ribosome-bound fractions (using a published RNA-seq dataset) (Data S8). Text adjacent to representative plots indicate KEGG terms related to the genes in each respective group (Data S9). Plot colours depict RNA (blue), RiboSeq (green) and Protein (red). (B) Compendium of encoded Arabidopsis proteasome subunits (Data S10). Genes displaying transcriptional (blue), protein (red) or both (purple) oscillations are shown, while white coloured nodes are those not detected in the study (Data S1-S5). (C) Diel transcript and protein profiles for select core and regulatory proteasome components. Relative transcript (blue) and protein (red) abundances for each gene are shown. Bold lines represent genes displaying a significant diel oscillation (Data S5). The line and shading in (A) and (C) are medians and min/max values of 3 replicates. Bold lines indicate significant rhythmicity. Highlighted genes include: RPN13 (REGULATORY PARTICLE NON-ATPASE 13; AT2G26590), SBP2 (SELENIUM-BINDING PROTEIN 2; AT4G14040), PAD1 (20S PROTEASOME ALPHA SUBUNIT PAD1; AT3G51260), RPE (D-RIBULOSE-5-PHOSPHATE-3-EPIMERASE; AT5G61410), G6PD5 (GLUCOSE-6-PHOSPHATE DEHYDROGENASE 5; AT3G27300).

One mechanism of post-translational regulation of protein levels that has previously characterized links to the plant circadian clock is proteasomal degradation(*30*). Targeted investigation of the proteasome found extensive transcriptional-related connections to the circadian clock that spanned all facets of the multi-subunit complex (Fig. 6B - 6C; fig. S9; Data S10). In order to ensure that our analysis of proteasome subunits was not biased due to differences in the protein half-lives of core and regulatory subunits, we also performed a correlation analysis of protein amplitude and peak ZT for core and regulatory proteasomal proteins with half-life data obtained from Li et al. (*44*) (fig. S10). Here, no clear correlation was observed, suggesting that the regulation of time-of-day abundance of these subunits is disconnected from their overall half-life. Diel transcriptional oscillations of the core proteasome subunits remained largely unaffected in the circadian clock mutants examined; however, their diel proteomic oscillations were completely abolished (Fig. 6B - 6C; Data S10). A more diverse set of diel transcript-protein responses was observed in the regulatory proteasomal subunits, including complete abolition of both diel proteomic and transcriptional oscillations of RPN3A in each of the clock mutants studied here (Fig. 6B; fig S9; Data S10). However, the function of these regulatory subunits is contingent on the core proteasome(*45*), reinforcing our hypothesis that precise timing of diel proteostasis is key to how the circadian clock genes connect diel rhythms of gene expression to functional protein-level outcomes in plants.

## CONCLUSIONS

In this study, we reveal that the plant circadian clock exerts far greater control over diel protein abundance than previously recognized, reshaping long-standing assumptions about primarily transcriptionally-driven circadian clock regulation in plants. Using a data-independent acquisition based mass spectrometry proteomics workflow, which is less prone to the biased detection of high abundant proteins, coupled with parallel transcriptomics, we found that more than one-fifth of the quantified Arabidopsis proteome oscillates under diel cycles. By simultaneously profiling a set of well characterized circadian clock mutants, we discovered that different clock components play a role in synchronizing the rhythmic proteome to a bimodal phase distribution. In particular, we found that the morning module, LHY / CCA1 plays a uniquely strong role in maintaining proteome-wide synchrony in diel oscillations. These findings establish proteome rhythmicity as a key output of the circadian clock network.

Studying the rhythmic proteome of wild-type plants revealed intriguing patterns where proteins involved in energy metabolism have a more temporally distributed expression pattern, while proteins involved in specialized metabolic pathways such as glucosinolate metabolism are much more temporally restricted. This was mirrored in our analysis of transcript-protein phase decoupling, suggesting again a role for post-transcriptional regulation in maintaining these distinct regulatory architectures. Observations such as the distinct timing of nuclear- and plastidial-encoded components of the plastid ribosome showcase how our data can assist plant scientists studying a wide-range of biological processes to discover, and study, how those processes are shaped by diel regulation. To help facilitate this, we have also established an ePlant module on the Bio-Analytic Resource (BAR; https://bar.utoronto.ca/∼dev/eplant/; fig S11) to accelerate community advancement.

Contrary to the proteome-level disruption observed in clock deficient plant lines, to our surprise, the transcriptome remained broadly rhythmic across genotypes, revealing a striking disconnect between rhythmic mRNA production and rhythmic protein accumulation under diel conditions. Our results suggest that while light-dependent processes significantly impact the transcriptome, the circadian clock acts at the protein-level to maintain rhythmicity in gene expression. Comparing proteomic, Ribo-Seq, and RNA-seq datasets demonstrated that only a small fraction of rhythmic proteins are controlled solely at the transcriptional level, whereas most are likely regulated through translational or post-translational mechanisms. This remains comparatively unexplored in plants relative to model non-photosynthetic systems.

The plant circadian clock has been canonically understood to impact downstream processes transcriptionally, due to its composition as a network of transcription factors that regulate the expression of their target genes by promoter-binding. On the contrary, our results call for a new understanding of circadian-related diel plant cell regulation in which the clock exerts far greater and wide-ranging control at the translational and post-translational level. By demonstrating pervasive clock control over proteostasis, translation, and protein-specific timing, we show how the circadian clock network orchestrates temporal compartmentalisation across plant metabolism, growth, and cellular maintenance. Future work in this direction should involve targeted analysis of proteasomal and ribosomal deficient plants as well as profiling of protein post-translational modifications in clock mutants. The recent optimisation of tools such as Bio-Orthogonal Non-Canonical Amino acid Tagging (BONCAT) for plant science (*46*) also suggests new possibilities to profile proteostasis across the day. Together with the data presented in this paper, this would allow for a deeper understanding of how the plant clock synchronizes the rhythmic proteome in a diel manner.

## METHODS

### Plant Growth and Harvesting

Wild-type *Arabidopsis thaliana* Col-0 (WT), *lhy-20 cca1-1* (*lhy cca1*; Gift from Jose Pruneda - University of California San Diego), *prr7-3 prr9-1* (*prr7 prr9*) (*47*), *prr5-11 prr7-11* (*prr5 prr7*) (*48*), *toc1-2* (*toc1*)(*49*), *elf4-101* (*elf4*)(*50*), *lux-4* (*lux*)(*51*) were sown in soil (Sunshine Mix #1) with four plants per pot, stratified for 3 days at 4 °C and then germinated under a 12h : 12h light : dark photoperiod with 120 μmol/m^2^*s fluorescent light at 22°C. At 24 days after germination, four whole rosettes (pre-bolting) were harvested per genotype, per time-point, per sample and frozen in liquid nitrogen. Hence, each biological replicate was a pool of four rosettes grown in the same pot. Samples were homogenized using metal beads using a Genogrinder (SPEX Inc.) and aliquoted under liquid nitrogen for downstream analysis.

### RNA-seq and transcriptome data analysis

Total RNA was isolated from WT, *lhy cca1*, *prr7 prr9* and *toc1* plants for subsequent analysis as previously described(*52*), without deviation (n = 4). Briefly, total RNA was isolated from rosette material using the Direct-zol-96 MagBead RNA kit (Zymo Research). RNA integrity was assessed on a TapeStation 4150 (Agilent) according to the manufacturer’s instructions. To enable cost-efficient transcriptome profiling, bulk RNA was processed using a miniaturized Smart-seq2–based workflow as previously described(*52*). cDNA was amplified for 23 PCR cycles, and the resulting products were cleaned using Agencourt Ampure XP beads (Beckman Coulter) at a 0.8× bead-to-sample ratio. cDNA size distribution was evaluated using a Fragment Analyzer (Advanced Analytical) with an NGS Fragment High Sensitivity Analysis Kit, and concentrations were determined using a Qubit High-Sensitivity dsDNA assay. Library construction used the Nextera XT DNA Library Preparation Kit (Illumina), following the standard workflow but with all reaction volumes reduced to 1/10 to enable automated handling on an Echo LabCyte liquid handler (Beckman). Libraries were cleaned with Agencourt Ampure XP beads and again assessed using the Fragment Analyzer. Libraries were combined in equimolar proportions, and the final pool concentration was measured with the Kapa Library Quantification Kit (Roche). Sequencing was performed on a NovaSeq 6000 platform generating 150 bp paired-end reads.

Subsequent data analysis strategies for acquired transcriptome data was also performed as previously described(*52*), again without deviation. All subsequent bioinformatics analyses are described below. MultiQC was used for initial quality control(*53*). Samples that passed QC were aligned using STAR, and gene-level counts were generated using *featureCounts* within Rsubread (*54*). Rlog from DESeq2(*55*) was used for stabilizing variance across samples and count normalization.

### Quantitative proteomics sample preparation, mass spectrometry acquisition and data analysis

Total protein was isolated from WT, *lhy cca1*, *prr7 prr9*, *prr5 prr7*, *toc1-2*, *elf4*, and *lux* for quantitative proteome analysis as previously described (n = 4), without deviation(*52*). Briefly, the homogenate was portioned into ∼100 mg aliquots under liquid N₂. Each aliquot was suspended in protein extraction buffer [50 mM HEPES-KOH (pH 8.0), 100 mM NaCl, 4% (w/v) SDS] at a 1:3 (w/v) tissue-to-buffer ratio and incubated on an Eppendorf ThermoMixer F2.0 at 95°C with shaking at 1000 rpm. Following extraction, samples were centrifuged at 20,000 xg for 10 min at room temperature, and clarified supernatants were transferred to fresh tubes.

Proteins were reduced with 10 mM dithiothreitol (D9779, Sigma-Aldrich) for 5 min at 95°C and subsequently alkylated with 55 mM iodoacetamide (I1149, Sigma-Aldrich) for 30 min at room temperature. Total proteome peptide samples were then generated on a KingFisher Apex automated magnetic particle processing system (Thermo Fisher Scientific). Digestion was carried out using sequencing-grade trypsin (V5113, Promega) prepared in 50 mM triethylammonium bicarbonate buffer (pH 8.5; T7408, Sigma-Aldrich). After digestion, samples were acidified to a final concentration of 0.5% (v/v) trifluoroacetic acid (A117, Thermo Fisher Scientific). Desalting was performed using an OT-2 automated liquid handler (Opentrons Labworks Inc.) fitted with Omix C18 pipette tips (A5700310K, Agilent). Peptides were dried and stored at −80°C until being resuspended in 3.0% (v/v) acetonitrile and 0.1% (v/v) formic acid for LC-MS/MS analysis.

Peptide samples (1 μg per injection) were analysed on a Fusion Lumos Orbitrap mass spectrometer (Thermo Fisher Scientific) operated in DIA mode. Chromatography was performed using an Easy-nLC 1200 system (LC140, Thermo Fisher Scientific) equipped with an Acclaim PepMap 100 C18 trap column (164750) and a 50-cm Easy-Spray PepMap C18 analytical column (ES903) maintained at 50°C. Peptides were separated at 300 nL/min using a solvent B gradient of 0.1% (v/v) formic acid in 80% (v/v) acetonitrile from 4 to 41% over 120 min. Ion source settings included a spray voltage of 2.3 kV, an ion transfer tube temperature of 300°C, and an RF lens of 40%. BoxCar DIA acquisition was carried out as described previously(*29*). MS1 data were collected using two multiplexed targeted SIM scans comprising 10 BoxCar windows each, covering m/z 350 to 1400 at a resolution of 120,000 (200 m/z) with normalized AGC targets of 100%. MS2 spectra were collected at 30,000 resolution across 28 overlapping windows (38.5 m/z width, 1 m/z overlap), using a dynamic injection time, AGC target of 2000%, a minimum of six points per peak, and HCD fragmentation energy of 27%.

Raw data were processed using Spectronaut ver. 19 (Biognosys AG) by searching Araport11 (https://www.araport.org/), using a single missed cleavage and an FDR setting of 1% for protein, peptide and PSM as previously described(*29*).

### Bioinformatic Analyses

*Diel rhythmicity analysis:* Rhythmicity analysis of proteome (log-transformed) and transcriptome (rlog transformed) data was conducted using the Discorythm R package(*32*). For proteome data, which is inherently more variable, we called proteins as rhythmic when they were detected as rhythmic by either the Cosinor(*56*) or JTK-Cycle(*57*) algorithms with a *q-value* of <0.2, similar to other multi-omics circadian studies(*39*, *58–60*). For this analysis, proteins with missing values in any replicate within the same genotype were excluded to allow the algorithms to estimate rhythmicity. Furthermore, we assessed the impact of more stringent *q-value* thresholds on our rhythmicity analysis and found that the differences in proteome rhythmicity in WT plants and mutants was maintained at more stringent thresholds as well (fig. S12). For transcriptomic data, transcripts were considered as diel rhythmic when they were detected as oscillating by both Cosinor and JTK-cycle with a *q-value* of <0.01.

*Plotting transcript and protein abundance:* To compare the pattern of transcript/protein abundance throughout the day, the gene expression and protein abundance depicted in Fig. 6 were min-max normalised using the equation 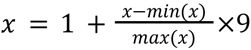. Where x’ is the normalised value, x is the original data value, and min(x) or max(x) are the minimum or maximum transcript/protein abundance of the gene across timepoints and replicates.

*Daily Expression Integral (DEI) analysis:* DEI of each protein/gene was calculated as the sum of the median abundance/expression level at each timepoint.

*Protein half-life calculation:* Protein half-lives were determined using first-order decay kinetics mathematical relationship from average protein degradation rate (K_d_) in Arabidopsis thaliana as reported(*44*). The conversion of degradation rate to half-life(*61*) was calculated using:

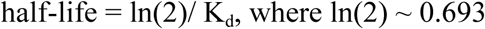

### Biological Pathway Analyses

To investigate the biological processes and functions of the genes identified from the transcriptomics and proteomics data, Gene Ontology (GO) and Kyoto Encyclopedia of Genes and Genomes (KEGG) pathway enrichment analyses were performed. Both were assessed using the same input AGI/gene data, in wild-type Columbia-0 (Col-0) background. Separate enrichment analyses were performed for the data sets obtained from transcriptomics and proteomics: transcripts, proteins, and transcript–protein pairs differentially expressed across the time course. For both GO and KEGG analyses, the background consisted of all AGIs identified within the corresponding experiment (i.e., AGIs significantly expressed in transcriptomics data set for transcript-level enrichment, AGIs with significantly different abundance from proteomics data set for protein-level enrichment, and the subset of genes/proteins commonly identified as considerably changing in both the data sets for transcript-protein pair enrichment). GO enrichment was performed using Ontologizer(*62*) with the parent-child union being selected as the analysis method. Benjamini-Hochberg FDR correction was performed and filtered for only the biological processes (BP) GO terms with adjusted *p-value* < 0.05. GO terms with population term size of 3-300 were used for transcript-protein pair enrichment analysis.

In contrast, the upper limit of the population term size was increased to 350 and 475 for transcripts and proteins, respectively, to account for the corresponding list size. KEGG pathway enrichment analysis was conducted using the *enrichKEGG* function from the *clusterProfiler* package (version 4.12.6) in R (version 4.4.1). The same background gene list and significance cutoff of adjusted *p-value* < 0.05 were used for the enrichment and filtering purposes. Figures were assembled using Affinity Designer ver. 2.0.

### Functional Association Network Analysis

Protein association networks of diel rhythmic proteins from WT, and genes with differential RNA-protein expression peak time were generated with Search Tool for the Retrieval of Interacting Genes/Proteins Database (STRING-DB v.12)(*41*). Interaction scores for edges were calculated based on sources including text mining, experiments, databases, co-expression, neighbourhood, gene fusion, and co-occurrence, with a cut-off score of 0.7. Single nodes without edges were filtered out, then functional enrichment analysis of the remaining nodes was performed against the genome with *stringApp* (v.2.1.1) in Cytoscape(*63*). Nodes with the same biological process were grouped together, with some manual adjustments based on protein descriptions. Subcellular localisation of proteins were annotated with Subcellular Localisation Database for Arabidopsis Proteins (SUBA)(*42*). For proteins with multiple localisation annotations, those annotated as ‘nucleus+cytosol’ were retained, the rest (12 out of 448 for WT, 4 out of 214 for RNA-protein) were manually annotated with representative subcellular localisations based on protein descriptions or published literatures.

### Community Visualization Tool

Normalized transcriptomic and proteomic data (see previous sections) were databased in a MySQL database on the BAR server. Rhythmicity testing results on protein abundance was performed for each protein group – genotype pair, only including statistically significant results. Outputs are included as waveforms fitted by the Cosinor and JTK-Cycle algorithms generated by Equation 1 below.

For all time points, x,

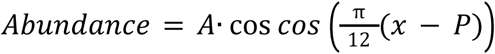

where A denotes amplitude, and P is the acrophase, both computed by Cosinor or JTK-Cycle. Javascript was the primary language used in the frontend responsible for dynamically generating the heatmaps in the new Combined Transcriptomics and Proteomics Viewer we developed for this project. Javascript was also used to implement user interactions within the viewer. The Circadian Proteomics Viewer’s user interface relies heavily upon the existing ZUI.ViewObject framework in ePlant(*64*). Chroma.js was used to easily compute colour gradients used in heatmap shading, and to allow for user-defined gradients. For review testing, the combined transcriptomics-proteomics viewer may be accessed at https://bar.utoronto.ca/∼dev/eplant/ .

## ACKNOWLEDGEMENTS

The authors would like to thank staff at the Alberta Proteomics and Mass Spectrometry Facility (APM; University of Alberta) and KU Leuven Genomics Core for assistance with data acquisition. As well, we thank G2V Optics Inc. (Edmonton, AB), for providing the LED lights used in the study. Lastly, we would like to thank the researchers who generously provided seeds for the plant lines used in the manuscript. This includes Drs. Jose Pruneda-Paz (University of California – San Diego; *lhy-20 cca1-1*), Eva Farre (Michigan State University; *prr7-3 prr9-11*), Philip Wigge (Leibniz IGZ & University of Potsdam; *lux-4*, *elf4-101*), Stacey Harmer (University of California Davis; *toc1-2*) and the Arabidopsis Biological Resource Center (ABRC; *prr5-11 prr7-11*, CS2107711).

## AUTHOR CONTRIBUTIONS

Conceptualization: DM, RGU

Methodology: DM, RGU, MCRG, SM, SL, DE Investigation: DM, MT, ZXL, QL, YZ, MCRG, SM, SL, DE

Visualization: DM, RGU, MT, ZXL, QL, YZ, SL, RL, WJL, WL, AP, NP

Funding acquisition: DM, RGU, DE, NP Project administration: RGU Supervision: DM, RGU, DE, NP

Writing – original draft: DM, RGU, MT, ZXL Writing – review & editing: DM, RGU, MT, ZXL

## FUNDING

**RGU**: Natural Sciences and Engineering Research Council of Canada (NSERC), Alberta Innovates Campus Alberta Small Business Engagement (AI-CASBE), Mathematics of Information Technology and Complex Systems (MITACS) Accelerate, Canadian Foundation for Innovation (CFI).

**DM**: Bijzonder Onderzoeksfonds start-up grant (#3E221118) by KU Leuven and Research Foundation-Flanders (FWO) project grant (#G081325N) to DM

**DE**: Biotechnology and Biological Sciences Research Council (BBSRC): BB/V006665/1

**NP**: Natural Sciences and Engineering Research Council of Canada (NSERC)

## DATA AVAILABILITY

All raw proteomics data can be found through the Proteomics IDEntifications Database (PRIDE; https://www.ebi.ac.uk/pride/) using the dataset identifier PXD071024. RNA sequencing data has been deposited in the European Nucleotide Archive (ENA) at EMBL-EBI (https://www.ebi.ac.uk/ena/browser/home) under accession number PRJEB103755. Additional processed Supplemental Data can be found on a Dryad Repository DOI:10.5061/dryad.h1893201s

---

**Fig. S1.**
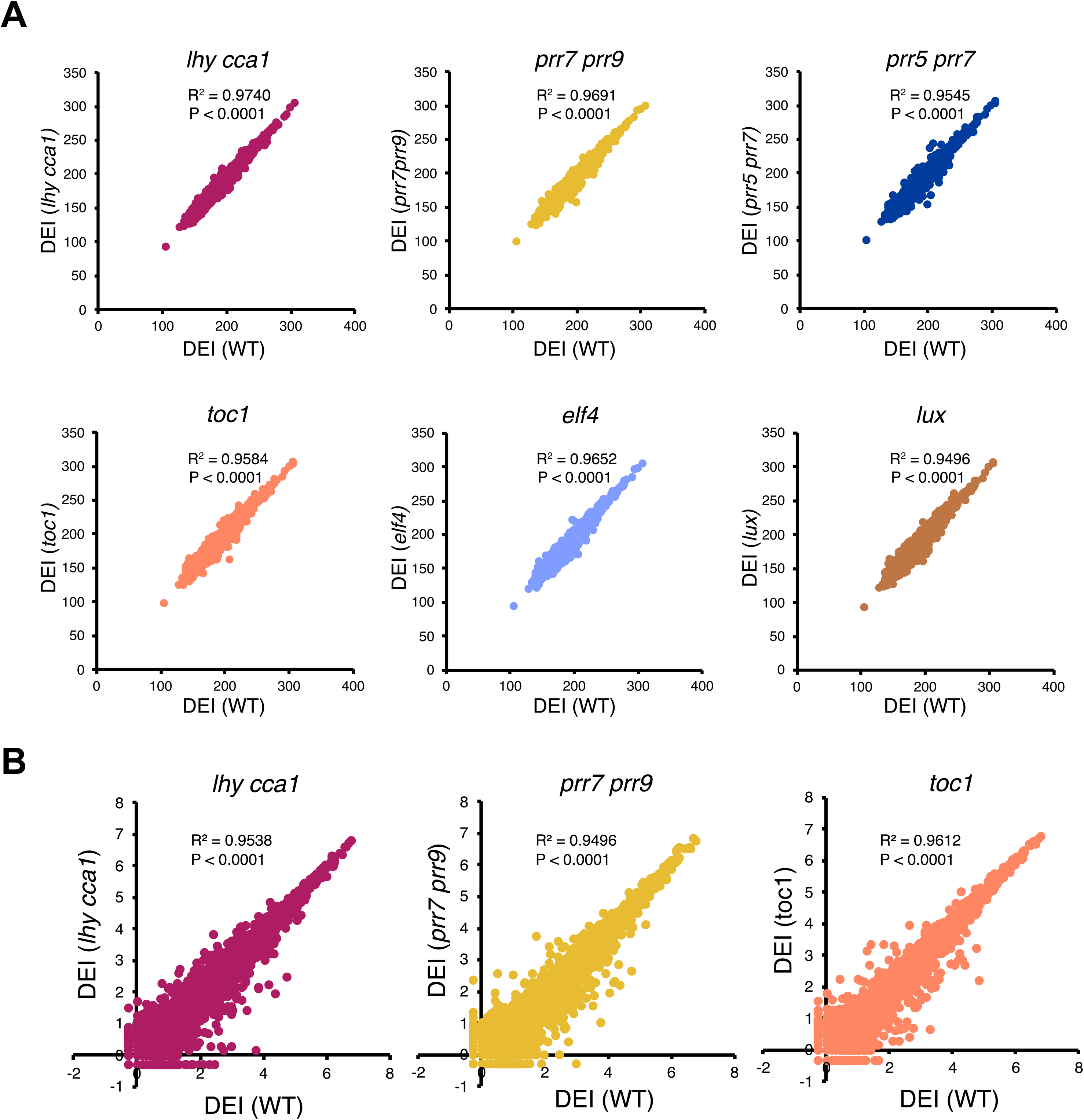
Correlation of the proteome/transcriptome daily expression integral (DEI) between WT and mutants. Correlation analysis of the DEI of proteome (A) or transcriptome (B) between WT and mutants. R^2^ and P values are calculated with Pearson correlation (details in methods).

**Fig. S2:**
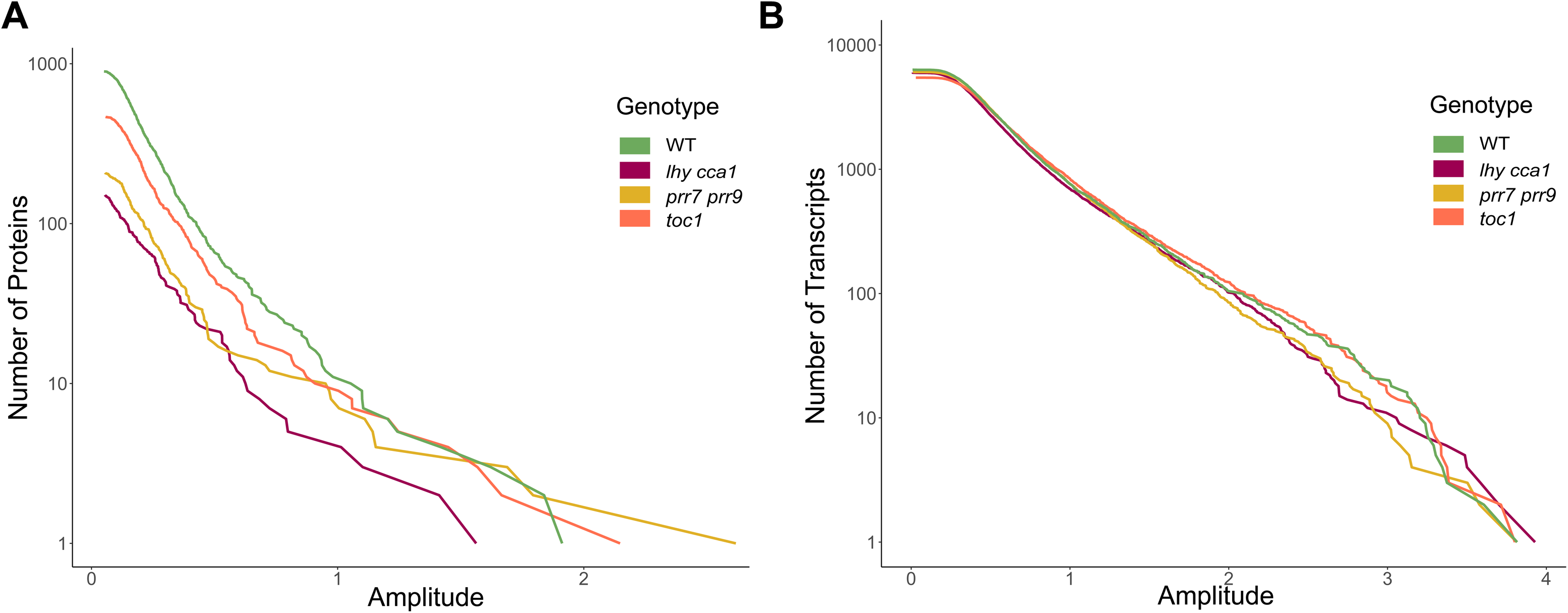
Distribution of rhythmic protein and transcript cosinor amplitudes by genotypes. (A) Cumulative number (Log_10_ scale) of rhythmic (A) proteins (Data S2) (B) transcripts (Data S4) across genotypes as a function of the amplitude of their rhythmic diel expression (estimated with Cosinor). Lines are colored by genotype.

**Fig. S3.**
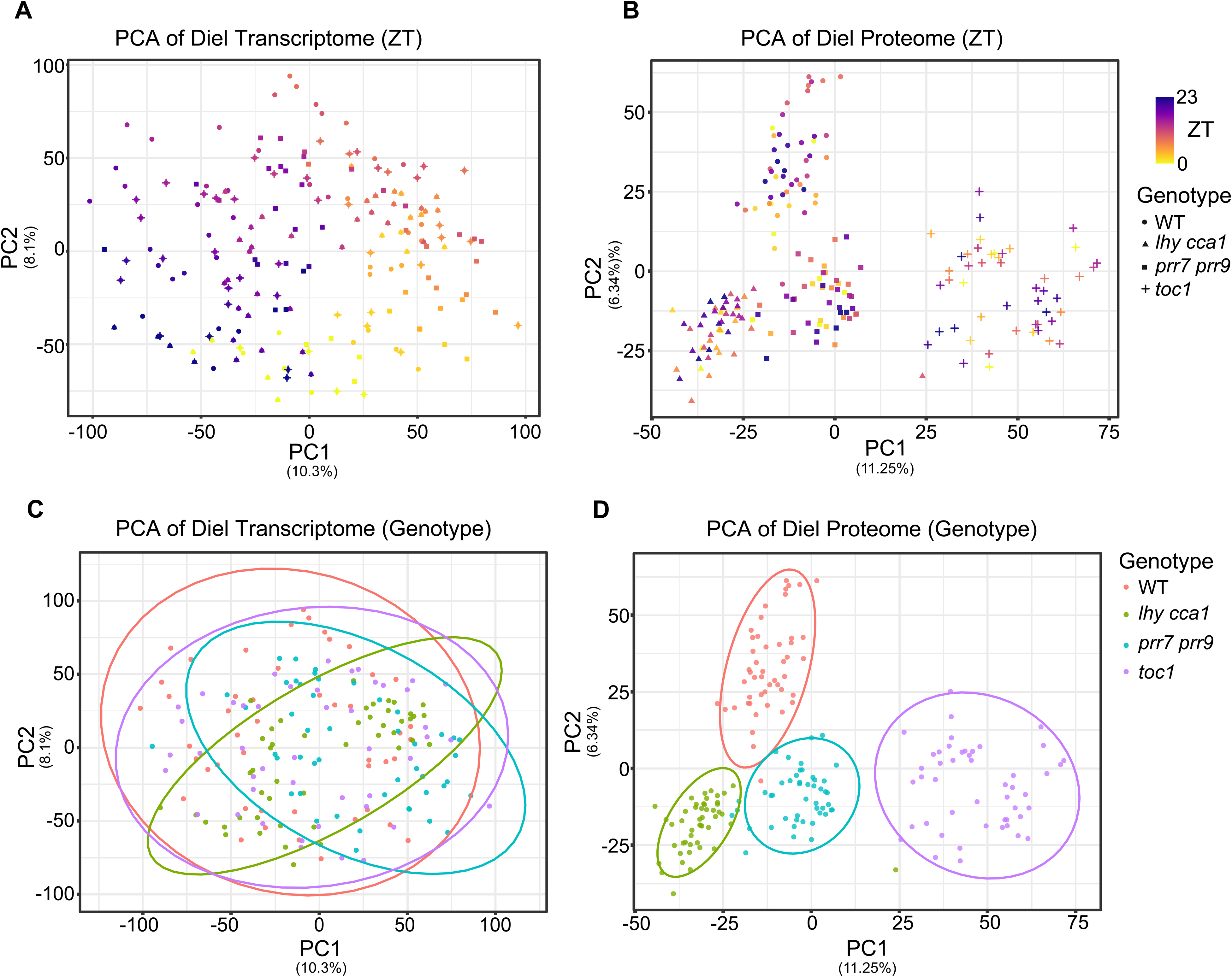
Principle Component Analysis (PCA) of the Diel Transcriptome and Proteome. PCA assessment of the diel transcriptome and proteome from WT, *lhycca1*, *prr7prr9* and *toc1*. The diel transcriptomes clustered by ZT, but not by genotype. Conversely, diel proteomes clustered predominantly by genotype. PCA is performed on all transcripts and proteins, not merely those classified as rhythmic.

**Fig. S4.**
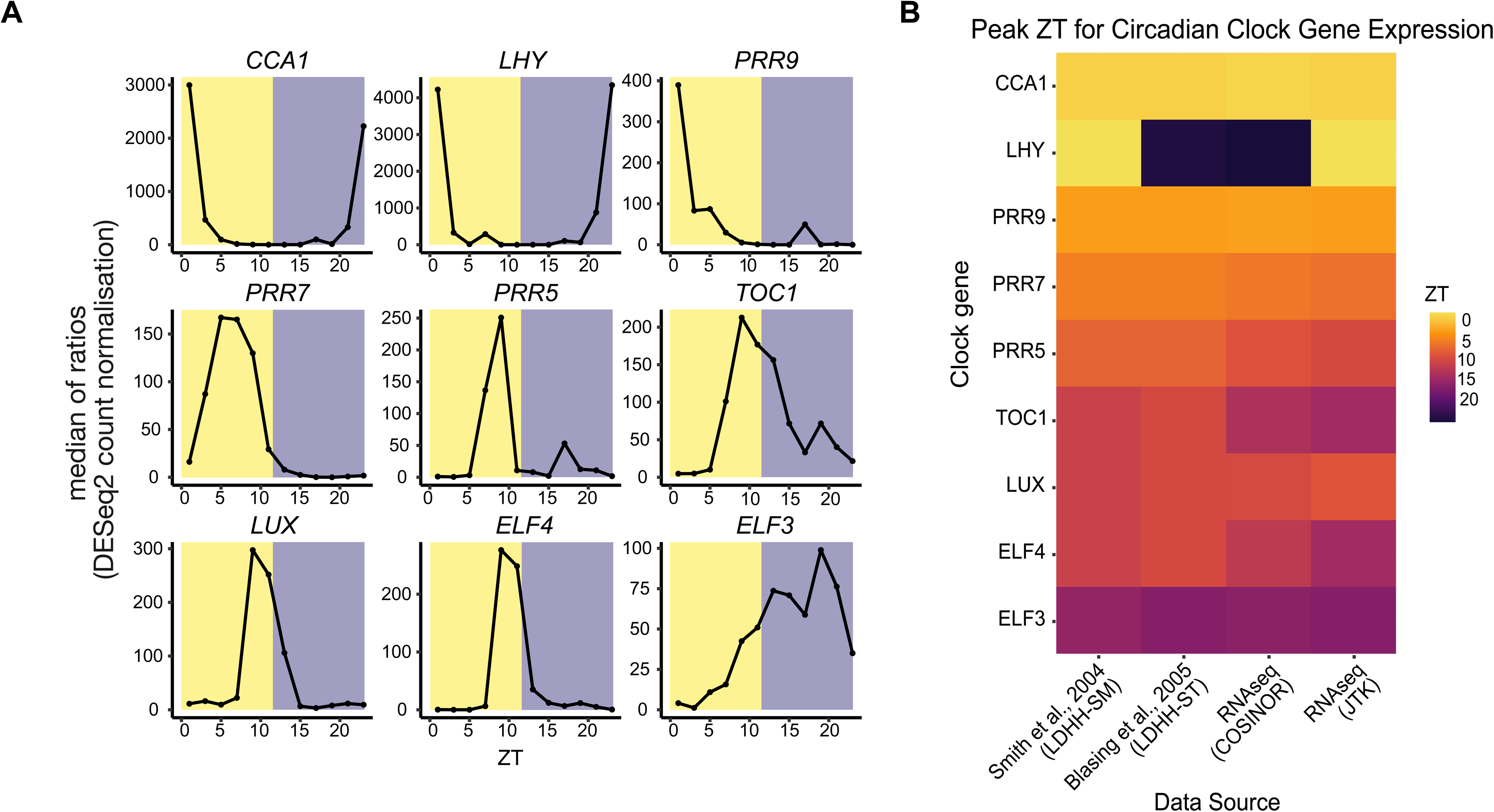
Benchmarking circadian clock genes’ transcript levels with previous datasets. Comparative peak expression analysis of clock genes within our dataset (RNAseq Cosinor & RNAseq JTK) against peak expression from Smith *et al.,* 2004 and Blasing *et al.,* 2005 obtained from DiurnalDB (http://diurnal.mocklerlab.org/).

**Fig. S5.**
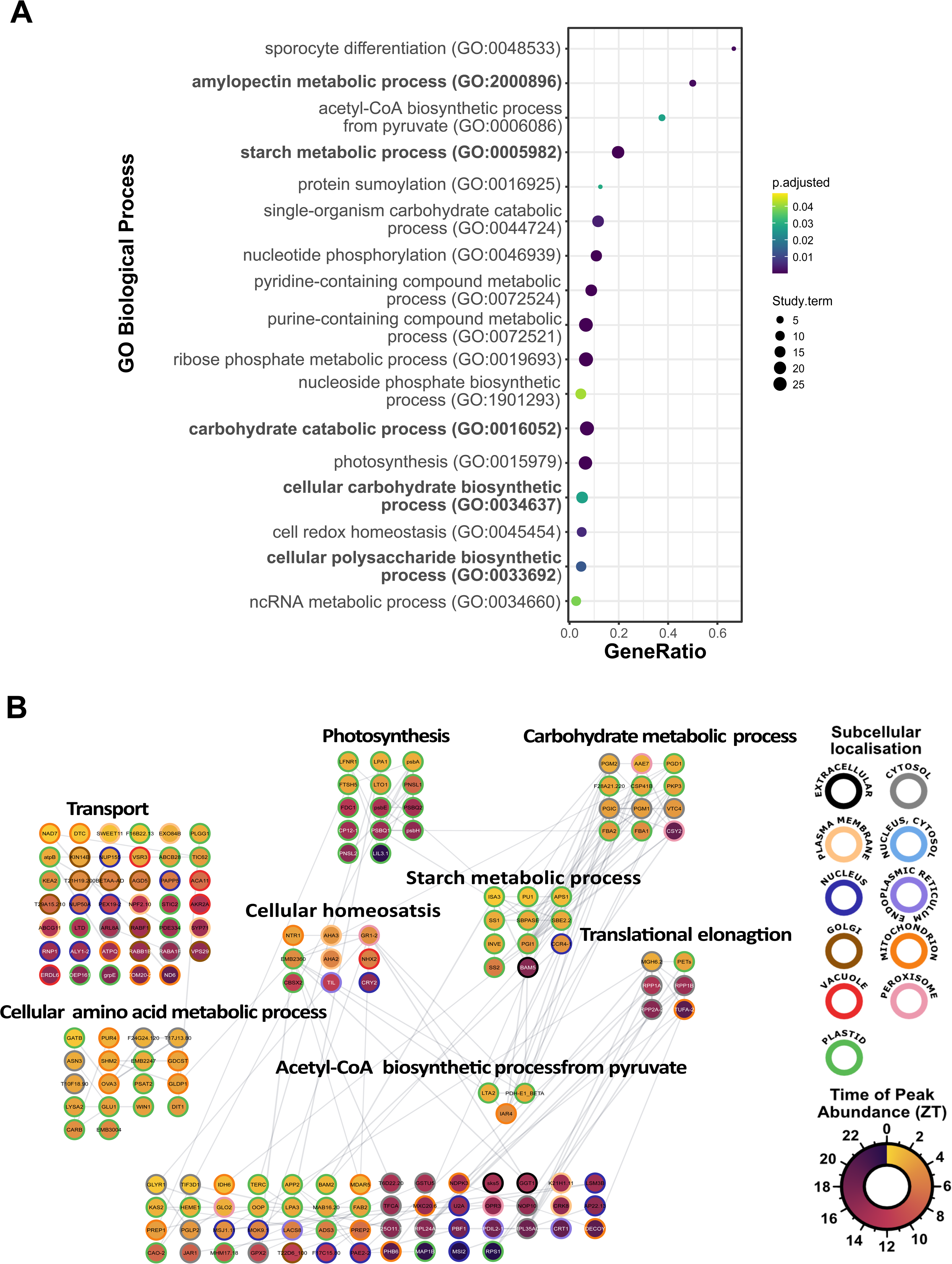
Genes with diel protein oscillations in *toc1* only. Upset plots in Figure 2C revealed that under diel conditions *toc1* had 265 genes with oscillating proteins that were exclusive to this genotype. (A) Gene ontology (GO) enrichment analysis (*BH p-value* < 0.05) and (B) functional association network of genes with diel oscillation in the *toc1* background exclusively (Cosinor; Data S1-S2 & S5).

**Fig. S6.**
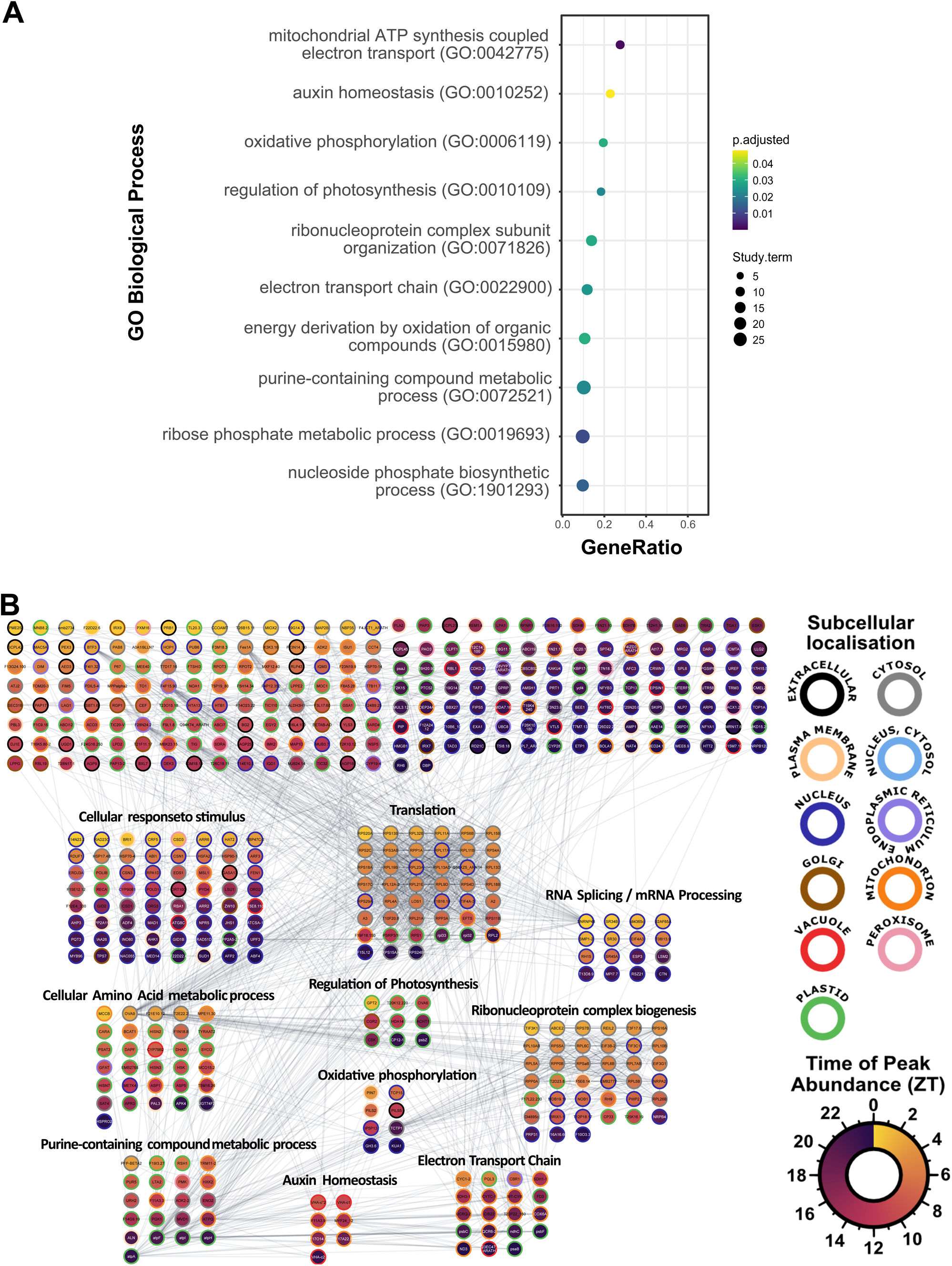
Genes with diel transcript oscillations in *lhy cca1* only. Upset plots in Figure 2C revealed that under diel conditions *lhy cca1* had 940 genes whose transcripts were diel rhythmic exclusively in this genotype. (A) Gene ontology (GO) enrichment analysis (*BH p-value* < 0.05) and (B) functional association network of genes with diel oscillation in the *lhy cca1* background exclusively (Cosinor; Data S4-S6).

**Fig. S7.**
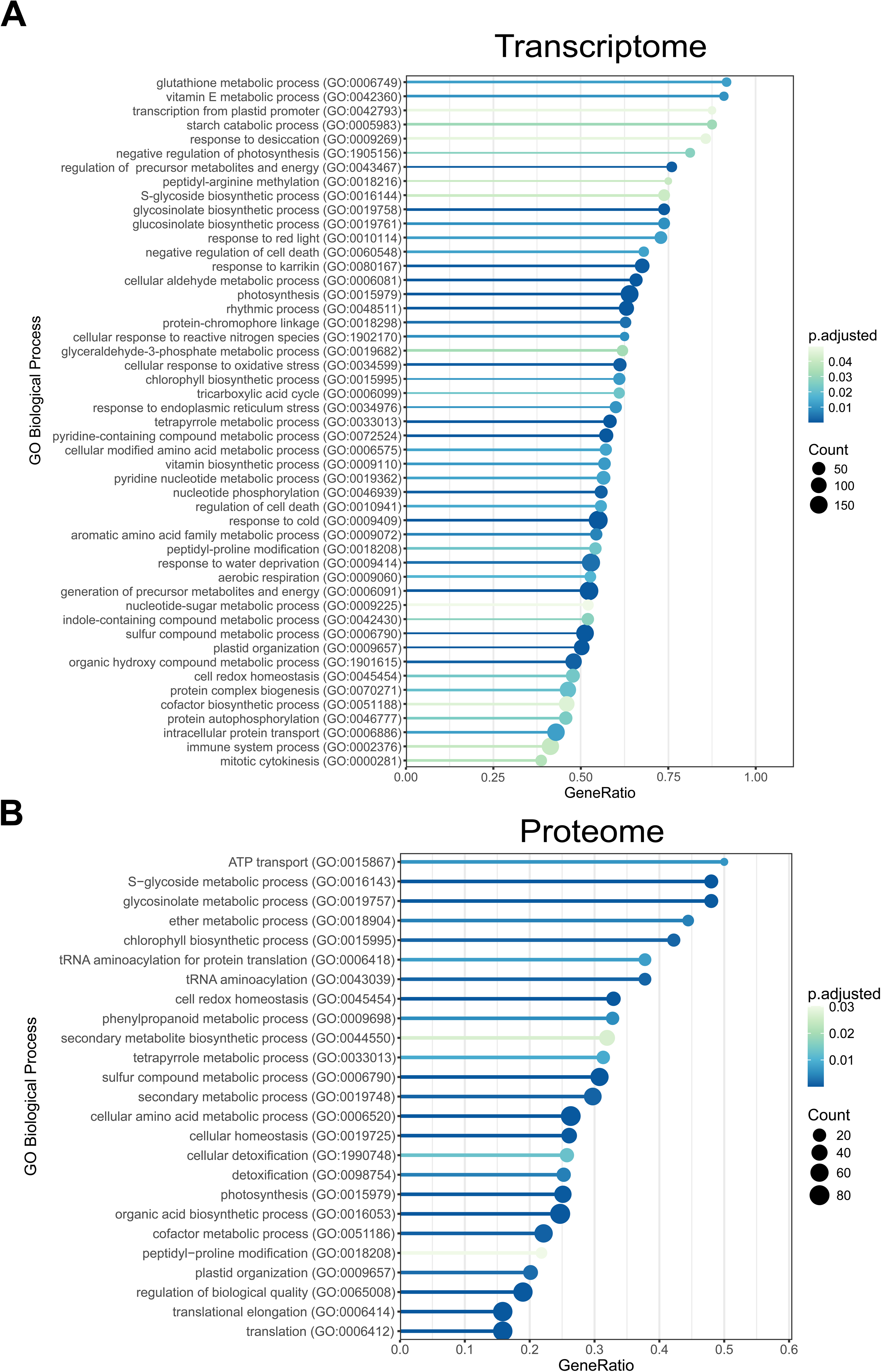
Gene ontology enrichment analysis of the diel transcriptome and proteome in WT Arabidopsis. The significantly oscillating transcriptome (A; Cosinor; Data S4-S5) and proteome (B; Cosinor; Data S2 & S5) were subjected to gene ontology enrichment analysis using The Ontologizer (http://ontologizer.de/; *BH p-value* < 0.05).

**Fig. S8.**
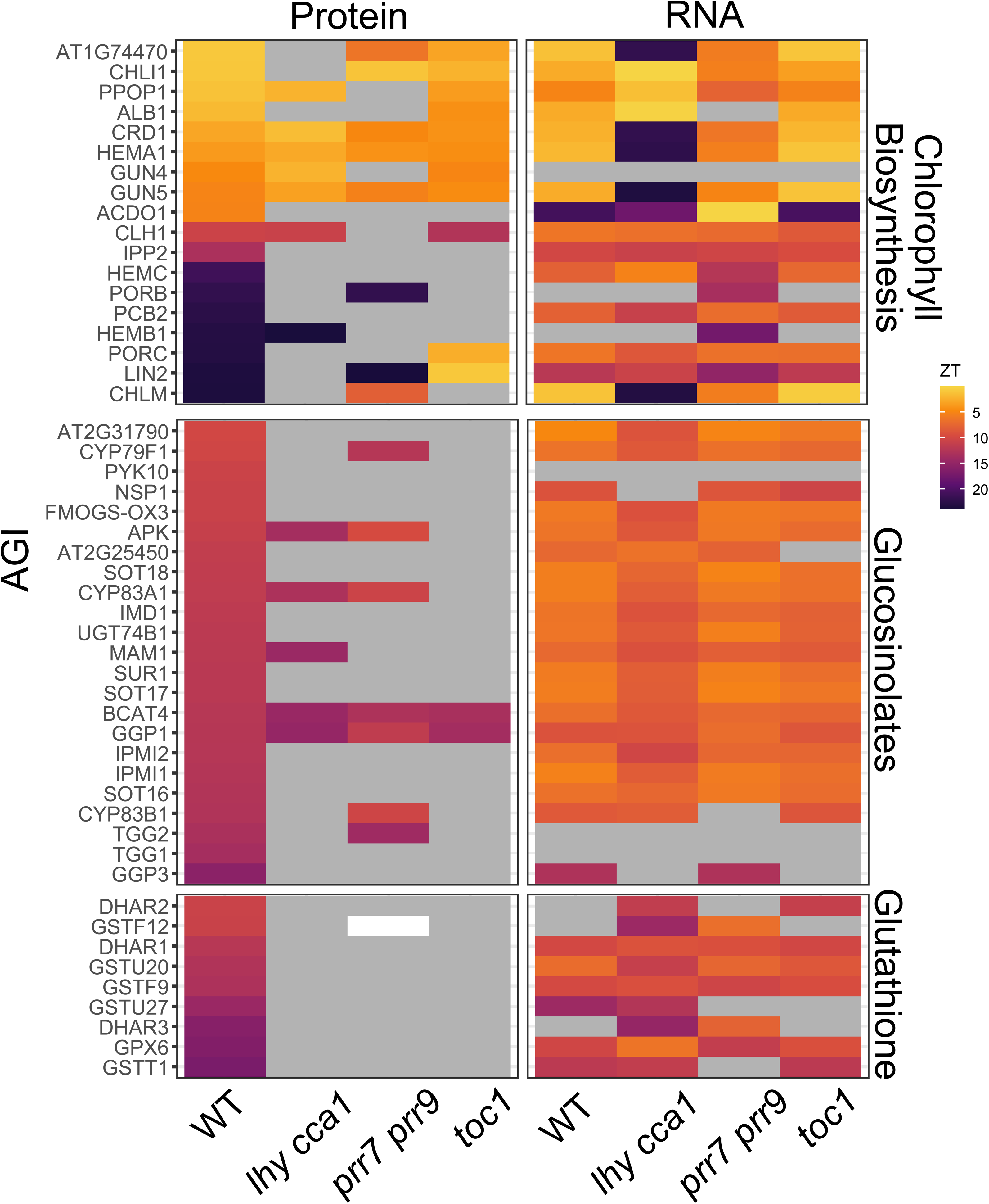
Biological processes governed by closely parallelized diel transcriptome and proteome changes. Genes involved in Chlorophyll, Glucosinolate and Glutathione Biosynthesis and Glutathione are extensively diel regulated at both the transcript- and protein-level. Similar to proteostasis-related proteins, loss of key circadian clock genes results in abolition of diel protein-level oscillations, while maintaining WT diel transcriptional oscillation patterns. Interestingly, morning phased chlorophyll biosynthesis genes maintain closely parallelized diel transcript and protein-levels even in the circadian clock mutants.

**Fig. S9.**
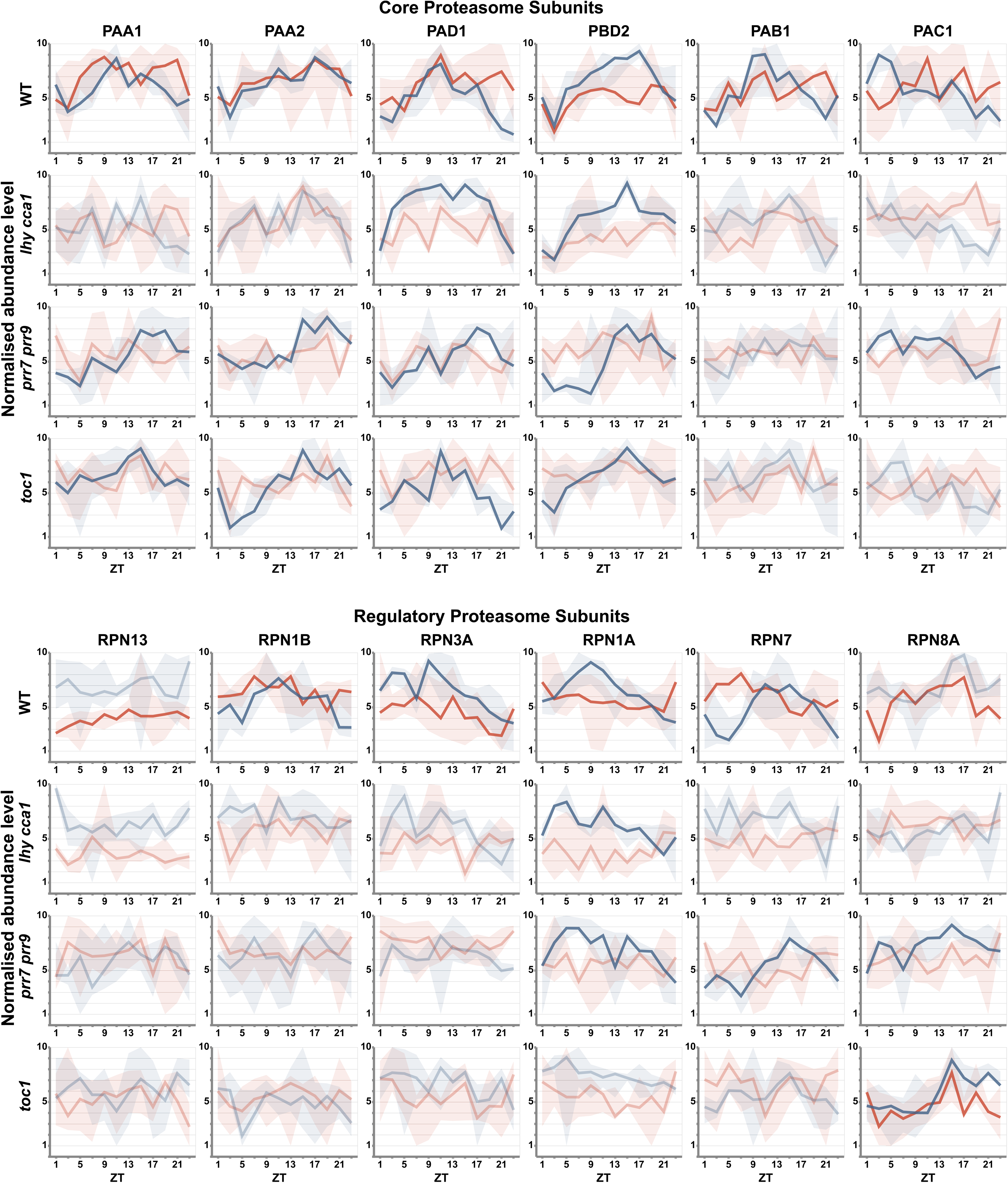
Diel transcript and protein profiles for all significantly oscillating core and regulatory proteasome components. Relative transcript (blue) and protein (red) abundances for each gene are plotted over the 24 h time-course. Bold lines represent genes displaying a significant diel oscillation in either transcript (Cosinor; Data S4-S5) or protein (Cosinor; Data S2 & S5).

**Fig. S10:**
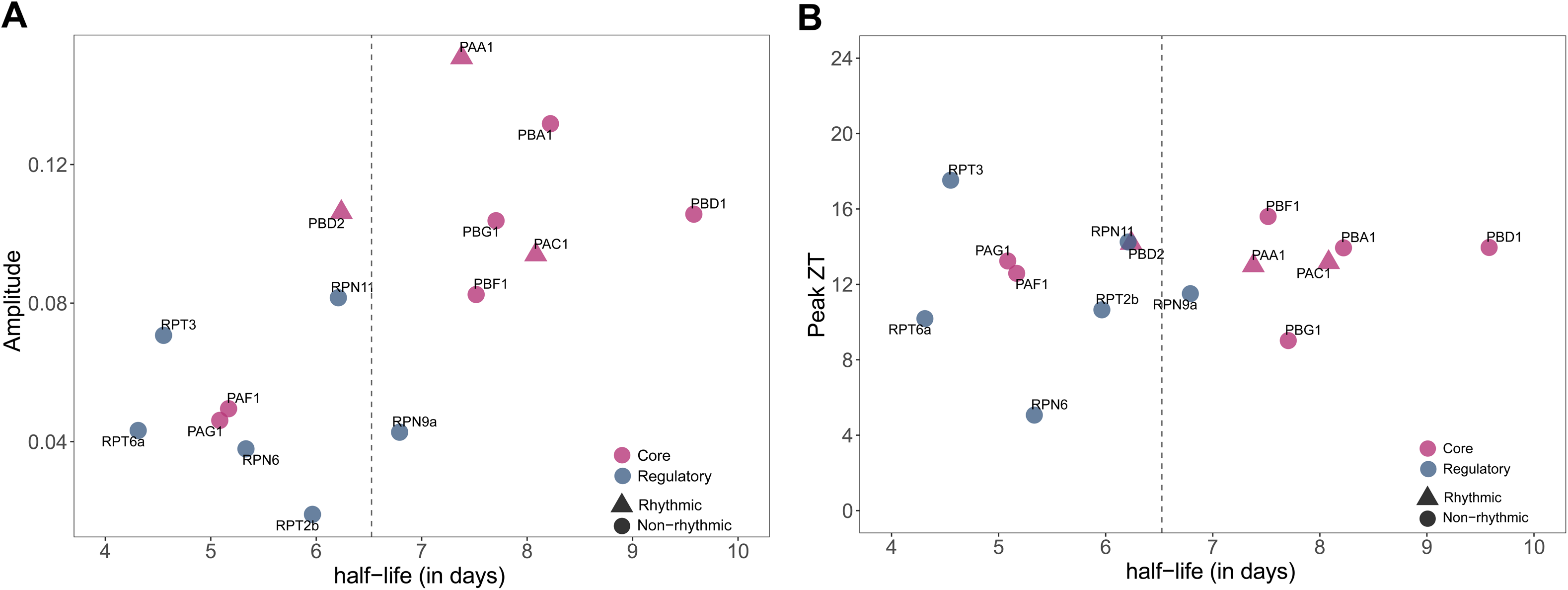
Proteasome subunit stability as depicted by protein half-life. Scatter plots showing the relationship between proteasome subunit half-lives (in days) and their changes in WT diel proteome (Data S6). Protein half-life, calculated from published protein degradation rate is plotted against (A) Cosinor Amplitude (Cosinor; Data S2) (B) Peak ZT (in hours) (Data S1). Each individual point represents a proteasome subunit, which are colored based on if they are core or regulatory unit. The point shapes denote the rhythmicity status based on rhythmicity analysis described in Methods. The vertical line represents the average half-life of all proteasome subunits that are included in the analysis.

**Fig. S11.**
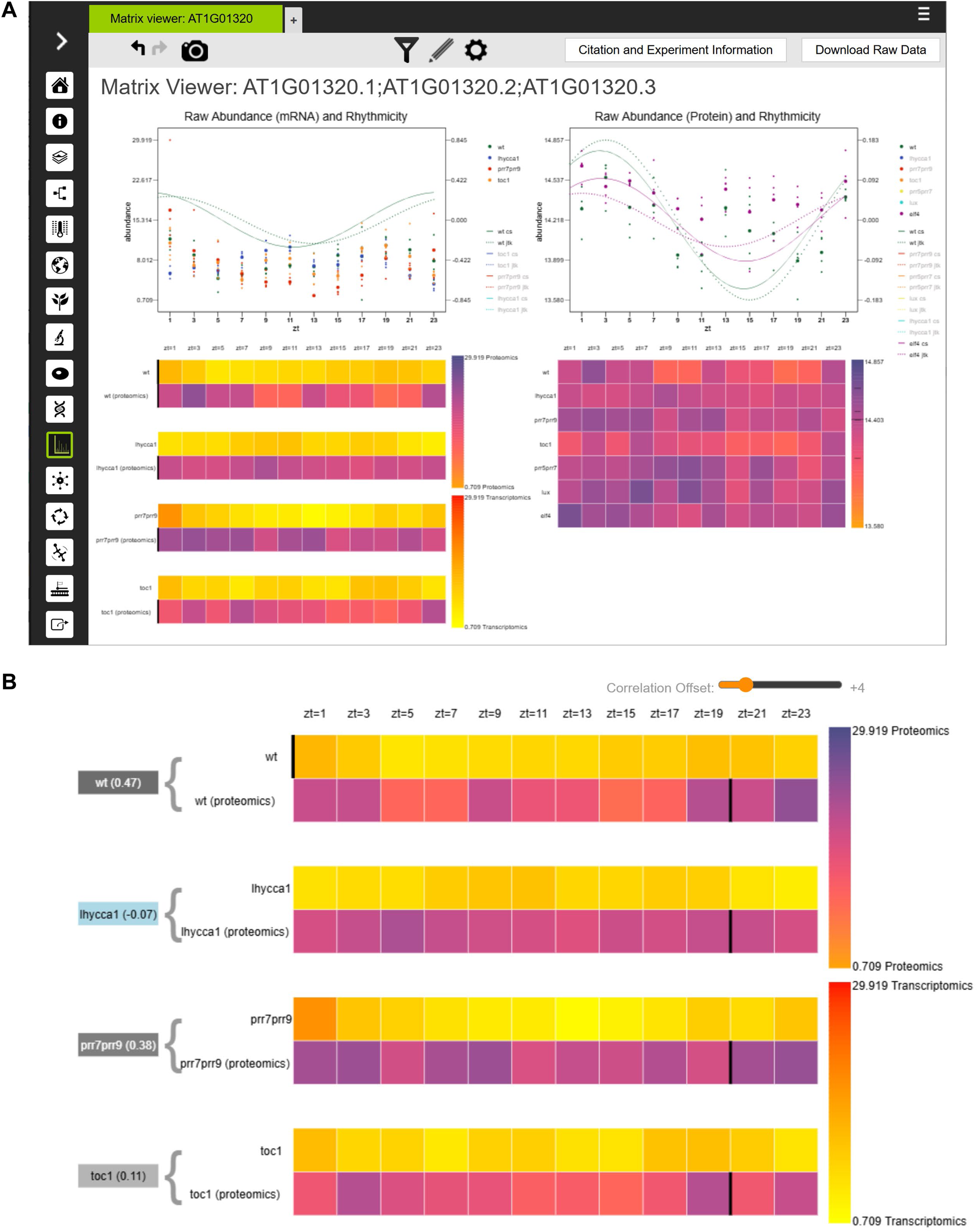
Bio-Analytic Resource ePlant module data integration and visualization. Screenshots of the newly developed combined transcriptomics-proteomics viewer for ePlant, for At1g01320, REC1, whose transcript/protein levels oscillate in a circadian manner, with a slight offset for the proteomics data in this case. (A) Screenshot of all parts of the viewer, with transcriptomics data presented in the top left graph, and proteomics data are in the right one. A stacked view (bottom left) in heatmap format of both data types allows similarities in response to be more easily discerned. The bottom right is a heatmap for just the proteomics data. (B) screenshot of the stacked viewer adjusted with a slider at the top right (“Correlation Offset”), activated by clicking the pencil icon, to allow the time offset for the calculation of the correlation between transcriptomics and proteomics data to be adjusted. The correlation score (as computed using the Pearson correlation coefficient) may be seen to the left of the stacked view. Here the offset has been adjusted by 4 hours, which results in an improved correlation for the WT samples between the transcriptomic and proteomics data from -0.17 to +0.47, whereby this improvement can be immediately seen by the colour scale behind the score, which is updated dynamically when the slider is moved – the darker the background colour, the higher the correlation.

**Fig. S12.**
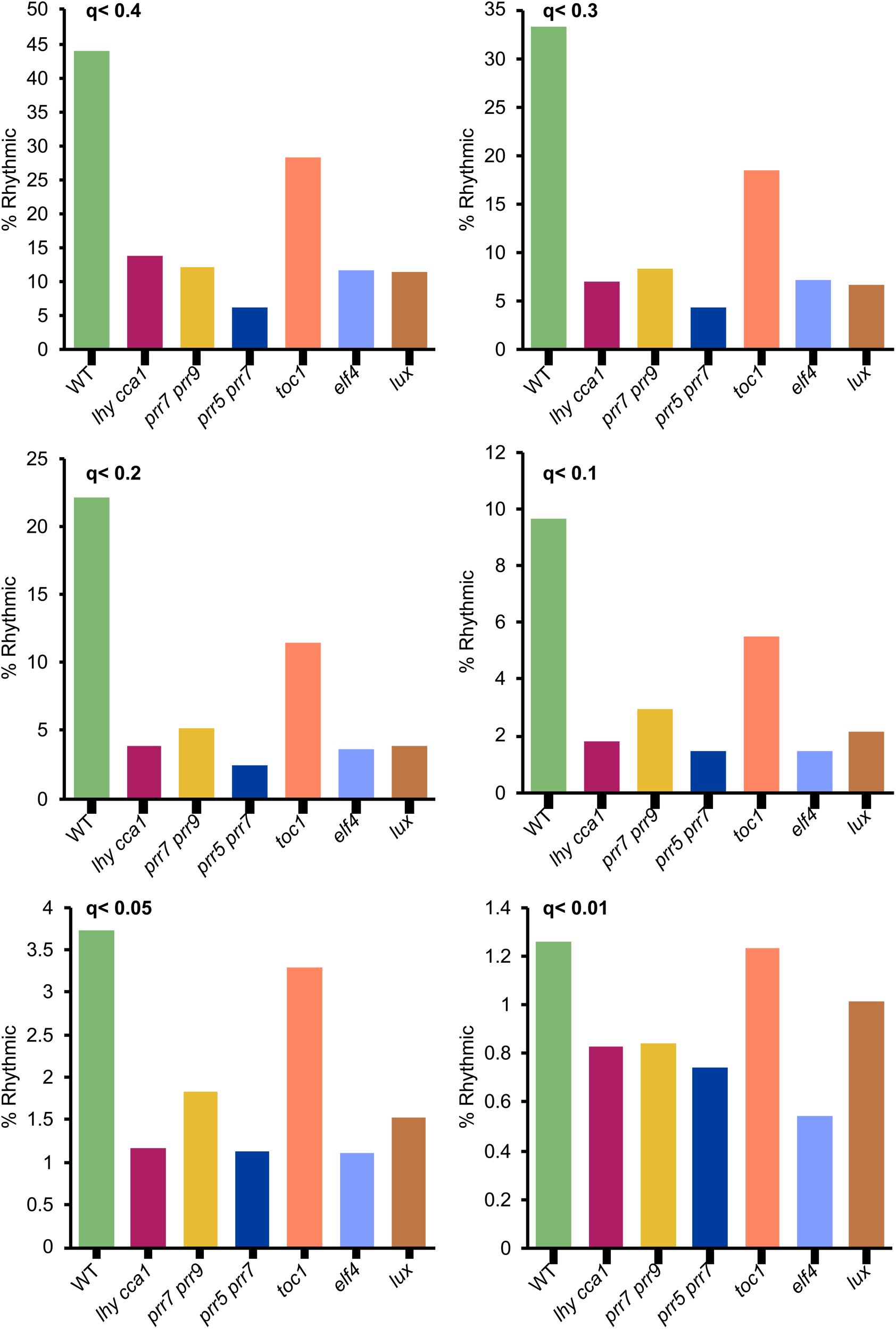
Effect of cut-off threshold on percent rhythmic proteins. Percent rhythmic proteins in WT and clock mutants, filtered with 6 different *q-values* for rhythmicity based on JTK-cycle/ Cosinor analysis as described in the Methods.

